# KaiC-like proteins improve stress resistance in environmental *Pseudomonas* species

**DOI:** 10.1101/2022.01.26.477890

**Authors:** Céline Terrettaz, Bruno Cabete, Johan Geiser, Martina Valentini, Diego Gonzalez

## Abstract

KaiC is the central cog of the circadian clock in Cyanobacteria. Close homologs of this protein are widespread among bacteria that are not known to have or need a circadian physiology. The function, interaction network, and mechanism of action of these KaiC homologs are still largely unknown. Here, we focus on KaiC-like proteins found in environmental *Pseudomonas* species. Using a bioinformatic approach, we describe the diversity and distribution of members of this protein family in the *Pseudomonas* genus and sketch, through comparative genomics, a conserved minimal interaction network comprising a histidine kinase and a response regulator. We then characterize experimentally the only KaiC homolog present in *Pseudomonas putida* KT2440 and *Pseudomonas protegens* CHA0. Through phenotypic assays and transcriptomics, we show that KaiC is involved in osmotic and oxidative stress resistance in *P. putida* and in sulfur uptake and alternative carbon source utilization in *P. protegens*. As expected, it physically interacts with its cognate histidine kinase and response regulator. Moreover, KaiC homologs are phosphorylated at one (*P. putida*) or two (*P. protegens*) sites and KaiC phosphorylation patterns change over time; however, in *Pseudomonas* species, changes in KaiC phosphorylation are driven by the age of the culture rather than by circadian cues as is the case in Cyanobacteria. In this study, through thorough bioinformatic and experimental analyses, we shed light onto the functional diversification and evolution of a unique protein family, diversely involved in bacterial rhythmic interactions with their environment. By so doing, we present a striking example of a protein whose general purpose is conserved in evolution, but whose molecular mechanics and participation in bacterial physiology can change dramatically across species.

## Introduction

KaiC is the ticking core of the circadian clock in the model Cyanobacterium *Synechococcus elongatus*. It is able, in interaction with two other proteins called KaiA and KaiB, to keep track of a 24-hour period in the absence of any external cue. The molecular functioning of this biological clock has been dissected in detail, demonstrating that all three Kai proteins are required to ensure persistence and entertainment of the system [1–3]. KaiC is an ATPase, whose active form is a hexameric; it has the capacity to autophosphorylate at, at least, two residues (a serine, S431, and a threonine, T432), when stimulated by KaiA, thereby generating four phosphorylation variants (S|T, S∼P|T, S∼P|T∼P, S|T∼P). The binding of KaiB to the hexamer prevents the stimulatory action of KaiA, inhibiting KaiC auto-kinase activity and stimulating its default auto-phosphatase activity; this in turn shifts the balance in the KaiA-KaiB competition towards KaiA and the cycle restarts. As a consequence, the relative abundance of KaiC phosphorylation variants oscillates in a 24-hour cycle *in vivo* and *in vitro*. The core oscillator indirectly drives circadian transcriptional programs. The main connection between the core and transcription goes through the response regulator RpaA, which is phosphorylated by the histidine kinase SasA when bound to the T∼P|S variant of the KaiC hexamer and dephosphorylates by histidine kinase CikA when bound to the T |S∼P variant of the KaiC hexamer. RpaA activates or represses transcription from two sets of promoters, leading to profound anti-correlated oscillations in the abundance of the two groups of transcripts [4]. Overall, more than 60% of *S. elongatus* transcripts are under circadian control [5].

Through bioinformatic analyses, it has been shown that KaiB and KaiC homologs were present in many archaeal and bacterial species throughout the phylogeny, whereas KaiA and most other members of the cyanobacterial input and output networks were restricted to a subset of Cyanobacteria [6–8]. Proteobacteria include a remarkable proportion of species encoding KaiB and KaiC modules or stand-alone KaiC homologs. Whether these KaiB and KaiC homologs are involved in circadian rhythms or not is not clear so far. The KaiB and KaiC homologs found in *Rhodopseudomonas palustris* (phototrophic alphaproteobacteria) are required for reliable circadian (hourglass-driven) nitrogen fixation and give a fitness benefit when light and dark alternate in 24-hour, but not one-hour, periods [9]. This strongly suggests that the KaiB and KaiC homologs are involved in circadian regulation in *R. palustris*, although their precise function and the connection with the input (environmental cues and relays) and output (nitrogen fixation) remain elusive. Interestingly, *cirA, cirB, cirC and cirD* genes, which encode four distant homologs of the cyanobacterial KaiC in the archaeon *Haloferax volcanii*, are induced by light and are also required for maintaining circadian rhythmic transcription [10]. By contrast, other reports highlight the involvement of KaiC-like proteins in stress resistance. In *Legionella pneumophila* (gammaproteobacteria), a KaiB and a KaiC homolog have been implicated in oxidative stress-response rather than circadian rhythms [7]. The transcripts of the two genes are regulated in a growth phase-dependent manner, by contrast with *S. elongatus*, in which the transcripts follow a circadian pattern. *Methylobacterium extorquens* (alphaproteobacteria) encodes two stand-alone KaiC homologs, KaiC1 and KaiC2, which are expressed in response to UV and heat or salt stress respectively [11]; they confer UV resistance in a temperature-dependent manner and have a variable impact on viability depending on the environmental conditions.

Here we explore the function, interaction network, and mechanism of action of KaiC homologs in environmental *Pseudomonas* species. The *Pseudomonas* genus is ubiquitous and highly adaptable to a great variety of environmental conditions; it has prodigious metabolic capacities, many strains being able to catabolize phenolic compounds and to synthesize secondary metabolites with potential medical or biotechnological uses. Some species are plant-adapted, like *Pseudomonas protegens* CHA0, which is well known as a biological control agent with fungicidal and insecticidal properties [12,13]. *Pseudomonas putida* KT2440 has been isolated from soil; it is able to degrade styrene and used as a biochemical synthesis platform in biotechnology [14–16]; like other *Pseudomonas* strains, it also shows light-induced phenotypes mediated by phytochromes and blue-light sensitive proteins [17–19]. These two species are representative of two of the main groups of environmental *Pseudomonas* and encode for a KaiC-like protein. As such, they are well suited to study the function of the KaiC protein family and its possible role in bacterial environmental adaptations.

## Methods

### Growth conditions

*P. putida* KT2440, *P. protegens* CHA0 and *E. coli* strains were routinely grown in LB broth (tryptone, 10 g/l; yeast extract 5 g/l; NaCl, 5 g/l; tryptone and yeast extract from Oxoid) or on LB supplemented with 1.5 % agar, except when indicated otherwise. M63 ((NH_4_)_2_SO_4_, 2 g/l; KH_2_PO_4_, 13.6 g/l; FeSO_4_•7H_2_O, 0.5 g/l; MgSO_4_, 1 mM; glucose 0.2 %) was used as a standard minimal medium. *E. coli* strains were grown at 37°C; *P. putida* and *P. protegens* strains were grown at 30°C; for liquid cultures, an agitation of 170 rpm was applied except when indicated otherwise.

When necessary, antibiotics were added to the media at the following concentrations (µg/ml) for (i) E. coli: kanamycin (50), tetracyclin (12), ampicillin (100), gentamicin (15) and (ii) Pseudomonas: kanamycin (50), tetracyclin (25 in LB, 12 in M63), ampicillin (100), gentamicin (30).

### Plasmids and strains construction

[Transformation conditions] Plasmid construction and amplification was done in *E. coli* DH5α-λpir chemocompetent cells prepared as the following: an overnight culture was diluted 100x in LB and grown until it reaches OD600=0.3-0.5; cells were chilled on ice and pelleted by cold centrifugation; pellets were resuspended in cold CaCl_2_ 0.1 M (1/2 of the culture volume) and incubated on ice for at least 30 min; cells were pelleted by cold centrifugation, resuspended in cold CaCl_2_ 0.1 M supplemented with 13 % glycerol, divided in 50 µl aliquots and immediately stored in −80° C. Transformations into *E. coli* chemocompetent cells were done using heat shock (42°C for 1 minute). Electroporations into P. putida and P. protegens were done at 25 µF, 200 Ohms and 2.5 kV after two washes of an overnight culture with a 15% glycerol, 1 mM MOPS buffer. Stocks of all strains were conserved at −80° C in 20% glycerol.

[Chromosomal *kaiC* deletion and FLAG *tagging*] To construct Δ*kaiC* deletion strains in *P. putida* KT2440 and *P. protegens* CHA0, the 400-600bps flanking the *kaiC* gene upstream and downstream were amplified by PCR separately using the Phusion DNA polymerase (New England Biolabs) and purified directly from the PCR mix or from an agarose gel (Gel and PCR clean-up kit, Macherey-Nagel); the two fragments were fused together thanks to a primer-encoded 20-30bps overlap during a second PCR amplification round and the product cloned into the pEMG suicide vector [20] using EcoRI and BamHI. The integrity of the inserted DNA fragment was verified by Sanger sequencing (Microsynth). The resulting pEMG plasmids were electroporated into the corresponding strains and transformants having integrated the plasmid were selected using kanamycin. To promote the excision of pEMG, which contains two I-SceI restriction sites, the pSW-2 plasmid, expressing the I-SceI restriction enzyme, was electroporated into the constructed strains and transformants were selected on gentamicin. Colonies were checked on plate for kanamycin sensitivity; the actual deletion of the *kaiC* coding sequence was verified using PCR. Finally, the resulting Δ*kaiC* strains were cured of pSW-2 through subculturing until gentamicin-sensitive colonies were obtained. A similar procedure was used to construct the *FLAG-kaiC* strains. Briefly, the 400-600bps upstream flanking region of the *kaiC* gene and the first 400-600bps of the *kaiC* coding sequence were amplified using overlapping primers, which added to the 5’ of the *kaiC* gene a DNA sequence encoding a double FLAG tag. The two fragments were fused during a second PCR round and the resulting product cloned into the pEMG plasmid. The expression of the construct was check by immunoblot using the procedure described below.

[Bacterial two-hybrid] Genes of interest were amplified by PCR from the genomic DNA of *P. putida* KT2440 and *P. protegens* CHA0 and cloned in both pKT25 and pUT18C plasmids using BamHI and EcoRI (New England Biolabs). The integrity of the cloned sequences was verified by Sanger sequencing (Microsynth). pKT25- and pUT18C-derived plasmids to test proteins interactions were co-transformed in *E. coli* BTH101; positive clones were selected on both kanamycin and ampicillin.

[Transcriptional fusion] The 198bps (*P. putida* KT2440) or 255bps (*P. protegens* CHA0) upstream of the predicted operon containing the *kaiC* gene were amplified by PCR using the Phusion DNA polymerase (New England Biolabs) and cloned between the EcoRI and HindIII sites of the pPROBE-TT’ [21], which encodes a promoterless green fluorescent protein (GFP). Plasmids were electroporated into *P. putida* KT2440, *P. protegens* CHA0, and their Δ*kaiC* mutants.

[STRP tagging] For production of the FLAG-STREP-tagged *P. putida* histidine kinase (PP_3835) and a subpart of it encoding specifically the PAS domain (amino acids 1-138) the corresponding coding sequence was amplified by PCR using the Phusion DNA polymerase (New England Biolabs) and cloned between the NdeI and BamHI sites, resulting in pET16b-HK_PP or pET16b-HK-PAS_PP plasmids, respectively. The obtained plasmids were transformed in *E. coli*, transformants were selected on ampicillin and the sequences integrity was verified by sequencing (Microsynth).

[His tagging] For production of His10Smt3-KaiC *P. putida* KT2440 (KaiC_PP) in *E. coli* BL21 (DE3), the *kaiC* gene was amplified from *P. putida* KT2440 genomic DNA and cloned into pET28b10xHis-Smt3 between HindIII site at the start codon and an XhoI site 3’ of stop codon. The resulting recombinant plasmid pSmt3-KaiC_PP encoded KaiC proteins with a N-terminal cleavable 10xHis-Smt3 tag. The constructs were confirmed by Sanger sequencing (Microsynth).

Plasmids, strains, and primers used in this study are listed in supplementary Table 1, 2, and 3 respectively. Strains built for the bacterial two-hybrid assays were not included in the strain list.

### Bioinformatics

1255 complete genomes within the *Pseudomonas* genus were downloaded from the NCBI FTP repository on 2020-06-16 and searched for KaiC and KaiB homologs. Briefly, hmm models were built, using *hmmbuild* (http://www.hmmer.org), based on alignments for the Conserved Protein Domain Family cd01124 (KaiC) and its parent cd01120 (RecA like NTPases), as well as cd02978 (KaiB) and its parent cd01659 (TRX superfamily) from NCBI CDD database (2020-07-14); the hmm for RadA (cd01121) was also added to the model since this protein is clearly distinct but closely related to KaiC; the resulting models were concatenated and indexed using *hmmpress* before being used against each individual predicted proteome using the *hmmscan* software (http://www.hmmer.org). 222 different KaiC homologs and no KaiB homolog were retrieved; the KaiC homologs were aligned using *muscle* [22] and a phylogenetic tree was built using *fasttree* with the -wag -gamma options [23]. The tree showed three broad phylogenetic subgroups of proteins, for each of which we made an individual hmm model using the aforementioned procedure; we ran *hmmscan* again against all the proteomes using all models combined and assigned each protein to a particular subgroup. *Pseudomonas* strains were classified into five main groups (*aeruginosa, fluorescens, putida, stutzeri, syringae*) based on a published global *Pseudomonas* tree [24]; when strains were missing in the published tree, they were classified on the basis of NCBI species assignment. Presence/absence full (Sfile 1) and summary table (Fig. 1B) were compiled from the hmm results based on species assignment. Logos were generated using online *Seq2Logo* with Shannon logo type and no clustering [25].

**Figure 1:**
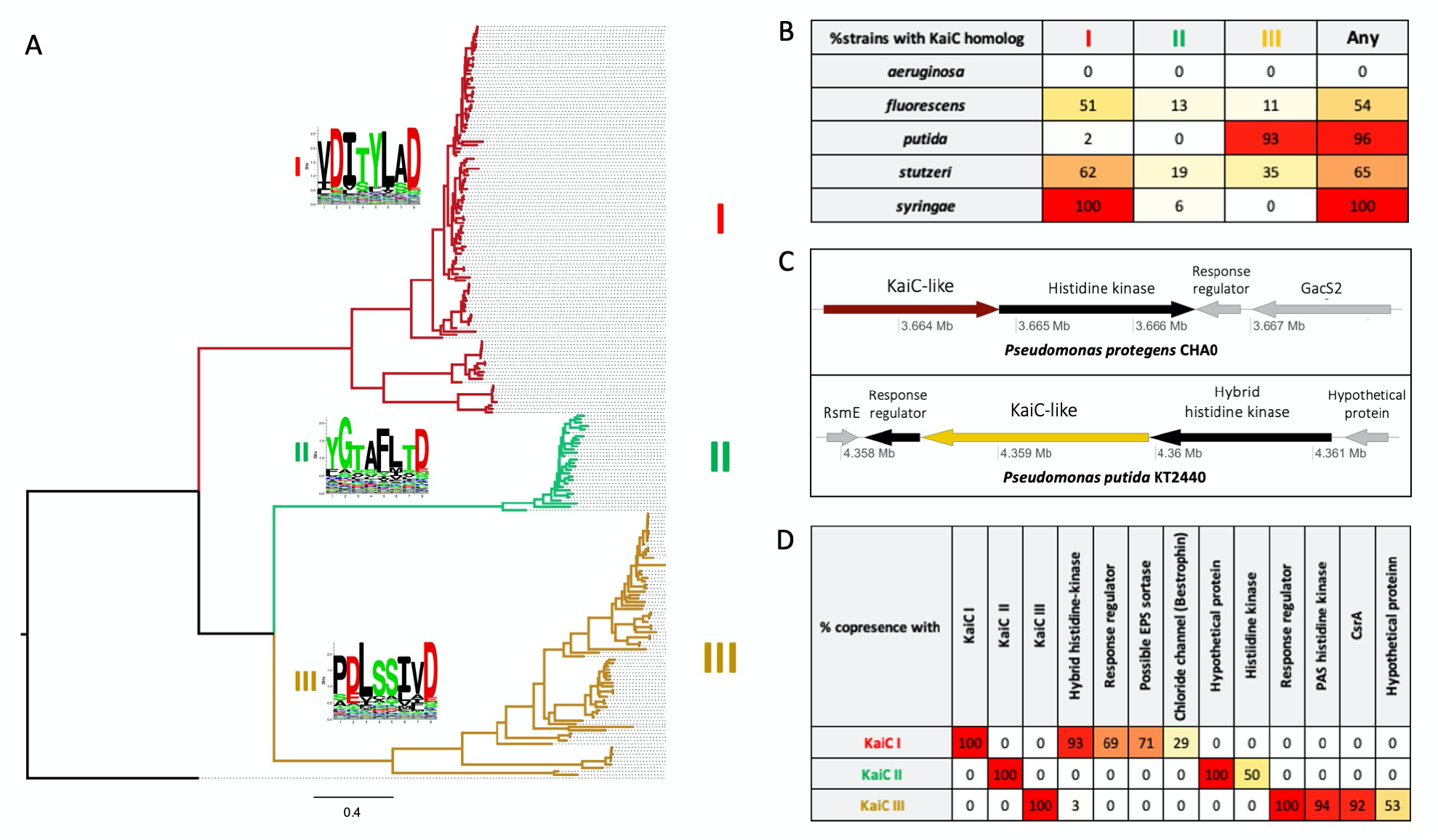
Three groups of KaiC-like proteins are found in environmental *Pseudomonas*. A. Phylogenetic tree of 222 KaiC-like proteins from *Pseudomonas* species rooted on the *Synecococcus elongatus* PCC_7942 KaiC. For each one of the three groups, a logo shows the representational bias of an 8-aminoacid region centered around the two residues that are phosphorylated in the Cyanobacterial KaiC (S^431^T^432^). B. Percent of *Pseudomonas* strains in each of five phylogenetic groups that encode a KaiC-like homolog belonging to groups I to III. KaiC-like proteins are found in virtually all “*putida*” and “*syringae*” strains and in more than half of the “*fluorescens*” and “*stutzeri*” strains. C. Genomic context around the *kaiC* gene in *P. protgens* CHA0 (group I KaiC, upper part) and *P. putida* KT2440 (group III KaiC, lower part); the genes coding the KaiC-like proteins are colored like the branches in panel A, genes in black are predicted to be co-transcribed, homologs of genes in grey are found in the neighborhood of KaiC-like homologs in other strains of the same group. D. Percent of genes coding for a KaiC-like protein, whose neighborhood (two genes upstream, two genes downstream) comprises genes coding for different protein models. A histidine kinase is virtually always present in the neighborhood of genes coding for KaiC-like proteins, while single domain response regulators are often present in groups I and III.

For each KaiC homolog found, we collected the protein sequences encoded by the two adjacent genes upstream and downstream, including the KaiC homolog, clustered them using *cd-hit* [26] with a 60% identity threshold, and used clusters with more than ten members to create an hmm model using *hmmbuild*. All *Pseudomonas* genomes encoding a KaiC homolog were searched with these models using *hmmscan* as described above. Summary tables were generated from the *hmmscan* output using *perl* scripts.

### RNA-sequencing

[KT2442 mid-stationary] Six cultures of *P. putida* KT2442 and its Δ*kaiC* mutant, inoculated to OD600 = 0.01 in LB, were grown for 24h before being pooled three by three and sampled (OD600 = 0.3, 1ml); pellets were immediately frozen at −80° C. For RNA extraction, samples were lysed with 1 ml TRIzol Reagent (ThermoFisher Scientific) at 65° C for 10 minutes; 200 µl of chloroform were added and, after centrifugation, 500 µl of the upper phase were taken. 300 µl of isopropanol were added to precipitate the RNA and after centrifugation and washing with 75 % EtOH, the RNA was resuspended in RNAse-free H_2_O. RNA was treated with DNase I (RQ1 RNase-free DNase, Promega) and purified with the *Quick-*RNA Fungal/Bacterial Microprep Kit (Zymo Research Corp.) including a DNase I on column treatment.

[KT2440 early stationary/late stationary and CHA0 late stationary] Triplicate precultures of *P. putida* KT2440, *P. protegens* CHA0 and their respective Δ*kaiC* mutants were grown overnight in LB. Culture triplicates were mixed, resuspended in 1.5 ml fresh LB and used to inoculate 4 cultures (500x dilution). After 12h (early) and 72h (late), cultures were sampled in triplicates (OD600 = 1.2, 1 ml). After centrifugation, cells were resuspended in 300 µl RNAlater (Sigma) and kept on ice for 30 min. The RNAlater was removed after centrifugation and the pellets were stored at −80 ° C.

For KT2440 early stationary phase samples, cells were lysed in TE buffer (pH=8) supplemented with 1 mg/ml of lysozyme and 10 µl of proteinase K (Qiagen) for 10 minutes at room temperature with a shaking of 500 rpm. The cells extracts were then mechanically lysed in a TissueLyser LT (Qiagen), 5 × 1 minute at maximal speed with 1 minute on ice in between. Samples were treated with the RNeasy mini-kit (Qiagen), following the kit instructions and including an on-column DNase treatment (RNase-free DNase set, Qiagen).

For KT2440 and CHA0 late stationary samples, cells were lysed in 1 ml TRIzol Reagent (ThermoFisher Scientific) at 65°C for 10 minutes; 200 µl of chloroform were added and, after mixing and centrifugation, 500 µl of the upper phase were mixed with an equal volume of 70% EtOH and loaded on a RNeasy mini-kit column (Qiagen). After an on-column DNase treatment (RNase-free DNase set, Qiagen), columns were washed and RNA eluted in RNase-free H_2_O according to the manual instructions.

RNA was quantified using Nanodrop2000 (ThermoFisher Scientific) and Qubit 2.0 fluorometer (Qubit RNA Broad Range Assay kit, ThermoFisher Scientific), checked for degradation on an agarose gel, and checked for possible DNA contamination by qPCR using SybrGreen (Rotor-Gene SYBR Green PCR kit, Qiagen) on a Rotor-Gene Q thermocycler (Qiagen). RNA-sequencing was carried out by Fasteris or Novogene on Illumina NovaSeq systems after in-house library construction. An average of 20 to 40 million 50bps pair-end reads (*P. putida*, mid-stationary, Fasteris) or 50 to 70 million 150bps pair-end reads (all other conditions, Novogene) were obtained per sample. Reads were filtered, trimmed of the first, low-quality, 5 bps (Fasteris) or 12 bps (Novogene), and searched for residual adapter sequences using *fastp* (v. 0.20.0) [27]. Remaining reads were mapped onto the *P. putida* KT2440 (NC_002947.4) or *P. protegens* CHA0 (NC_021237.1) genomes using *bowtie2* (v. 2.3.4.3) [28]. Reads were counted using *featureCounts* (v. 2.0.0) [29] with default count parameters. Analysis was performed in R (v. 4.0.0) [30] using the *DESeq2* (v. 1.28.1) package [31]. One sample was removed among the wild-type *P. protegens* CHA0 72h samples because of an anomaly in count dispersion. In the case of *P. putida* KT2440 72h, an additional parameter was added to the model to account for KaiC-independent consistent between-sample variations observed during data examination.

### KaiC promoter activity

Reporter strains carrying kaiC-transcriptional fusion were grown overnight in LB, washed and resuspended in fresh medium. Cultures were normalized to OD600 = 0.01 and loaded in quadruplicates in a 96-well plate. Optical density (OD600) and fluorescence (485/528 nm) were measured in a Biotek Cytation5 with moderate linear continuous agitation.

### Oxidative and osmotic stress

To test oxidative stress, strains of interest were grown overnight in LB. Cultures were vortexed for 30 sec, centrifuged and resuspended in fresh LB. OD600 was measured and adjusted to 0.002. Two-times concentrated H_2_O_2_ solutions in LB were prepared from a 100 mM stock. The cell suspension and the 2x H2O2 solutions were mixed 50:50; 6 replicates of 200 µl were loaded in a 96-wells plate and OD600 was recorded with a Varioskan Flash (Thermofisher) for up to 96 hours. The assay was also performed in M63 medium with a similar procedure, except that an intermediate preculture in M63 was done (500 x dilution of the washed LB culture).

To test osmotic stress, strains of interest were grown overnight in LB. After washing, cultures were adjusted to OD600 = 1, diluted 500x in M63 and grown for 24h. For the assay, cell density was adjusted to OD600 = 0.02. Two-times concentrated NaCl solutions were prepared from a 20 % stock, in M63. The cell suspension and the 2x NaCl solutions were mixed 50:50; 4 replicates of 200 µl were loaded in 96-wells plate and OD600 was recorded with a Biotek Cytation5 with moderate linear continuous agitation for up to 96 hours.

### SDS-PAGE, Phos-tag™ SDS-PAGE and immunoblotting

Over-night cultures of *P. putida* KT2440, *P. protegens* CHA0, and their Δ*kaiC* mutants were diluted to OD600=0.01 in LB and grown at 30°C with agitation for 6 hours (exponential phase), 12 hours (early stationary phase) or 24 hours (mid-stationary phase). Cells equivalent to 1 ml OD600=0.3 were collected; the pellet was resuspended into 300 µl of SDS-PAGE lysis/loading buffer (Wako Chemicals). 3 µl of sample were run on a 7.5 % SDS-PAGE or 7.5 % Zn^2+^-Phos-tag SDS-PAGE (SuperSep Phos-tag™, Wako Chemicals) in Tris-MOPS running buffer (Wako Chemicals guidebook) for 2h at 100V using a Mini-PROTEAN^®^ Tetra system (Bio-Rad Laboratories). After migration, the Phos-tag SDS-PAGE was quickly rinsed with deionized water, incubated in transfer buffer (Tris-base, 0.025 M; glycine, 0.19 M; ethanol 20 %; SDS 0.05 %) supplemented with 10 mM EDTA for 10 minutes to clear it from metallic ions and finally immersed in transfer buffer without EDTA. Proteins were then transferred onto a nitrocellulose membrane (Amersham™ Protran™ 0.2 µm NC, GE Healthcare Life science) using a standard blotting procedure (80V for 2h, in cold conditions, with stirring). Membranes were colored with amido-black (isopropanol 15 % (v/v), glacial acetic acid 10 % (v/v), amido black 0.1 % (w/v) in MilliQ water) to ascertain the equivalence of protein content between lanes. Membranes were then blocked in TTBS (Tris-base 0.05 M; NaCl 0.15 M; pH8 supplemented with Tween 20 0.1 %) supplemented with 5 % skimmed milk powder, incubated in TTBS supplemented with 2.5 % of skimmed-milk powder and a 1:10000 dilution of anti-FLAG antibodies (Sigma, F3165) for one hour and, after several TTBS washes, incubated with 1:10000 dilution of HRP-conjugated anti-mouse antibodies for one hour. After several washes in TTBS, luminescence was detected on an Amersham Imager 600 (GE Healthcare Life science) instrument after addition of freshly prepared ECL reagent (Tris-HCl pH8.5, 0.1 M; coumaric acid, 0.198 mM; luminol, 1.25 mM; 3 µl of 3 % H_2_O_2_ /ml of ECL reagent). Pictures were contrasted in *ImageJ* [32]; the same settings were applied to the whole image.

### Bacterial two-hybrid and β-galactosidase assay

Overnight cultures of *E. coli* BTH101 carrying the pKT25 and pUT18C plasmid derivatives of interest were washed with and resuspend in a NaCl 0.9 % solution. 5 µl of this cell suspension were spotted in triplicate on McConkey agar (Biolife) plates supplemented with 1 % maltose, 1 mM IPTG, kanamycin, and ampicillin. Pictures were acquired after 24h and 48h of growth at 30° C.

In parallel, 5 µl of the cell suspension were spotted in triplicates on LB agar plates, supplemented with 1 mM IPTG, kanamycin and ampicillin. After 24h of growth at 30° C, spots were collected and resuspended in 1 ml of a NaCl 0.9 % solution. OD600 was measured in a Varioskan Flash (Thermo Scientific). The cell suspension was lysed by addition of 40 µl of chloroform, vortexing and incubation for 15 minutes at 37° C. 20 µl of cell extract was mixed with 180 µl of Z-buffer (Na_2_HPO_4_, 0.06 M; NaH_2_PO_4_, 0.04 M; KCl, 0.01 M and MgSO_4_, 0.001 M) supplemented with 3.5 ml/l of β-mercaptoethanol. β-galactosidase assay was initiated by the addition of 40 µl of ortho-Nitrophenyl-β-galactoside (ONPG, 4 mg/ml in Z-buffer) and stopped after an appropriate reaction time with the addition of 100 ul of Na_2_CO_3_, 1 M. OD420 was measured in a Varioskan Flash (Thermo Scientific) and reaction times were recorded for data treatment.

Miller units were calculated as follows: (1000 × [OD420](/ ([Time, min.] × [volume, ml] × [OD600]) [33].

### Protein purification

The pSmt3-KaiC_PP plasmid was transformed into *E. coli* BL21(DE3). Cultures (500 ml) derived from single transformants were grown at 37°C in LB medium containing kanamycin until the OD600 reached 0.6. For each culture, protein expression was induced with 0.2 mM IPTG and 2% (v/v) ethanol and incubation was continued for 20 h at 17°C. Cells were harvested by centrifugation and stored at −80°C. All subsequent procedures were performed at 4°C. Thawed bacteria were resuspended in 25 ml of buffer A (50 mM Tris-HCl, pH 8.0, 200 mM NaCl, 10% glycerol) and supplemented with one tablet of protease inhibitor cocktail (Roche). The suspension was adjusted to 0.1 mg/ml lysozyme (Sigma) and incubated on ice for 30 min. Imidazole (Sigma) was added to a final concentration of 10 mM and the lysate was sonicated to reduce viscosity. Insoluble material was removed by centrifugation. The soluble extracts were mixed for 30 min with 3 ml of Ni2+-NTA-agarose (Qiagen) that had been equilibrated with buffer A containing 10 mM imidazole. The resins were recovered by centrifugation, resuspended in buffer A with 10 mM imidazole, and poured into columns. The column was washed with 15 ml aliquots of 20 mM imidazole in buffer A and then eluted stepwise with 5 ml aliquots of buffer A containing 50, 100, 250, and 500 mM imidazole, respectively. The 250 mM imidazole eluate fraction of each protein was dialyzed O/N in buffer A at 4°C and concentrated using Amicon® Ultra-4 centrifugal filters (Merk).

The pET16b-HK_PP or pET16b-HK-PAS_PP plasmids were transformed into *E. coli* BL21(DE3). Cultures (500 ml) derived from single transformants were grown at 37°C in LB medium containing kanamycin until the OD600 reached 0.6. Protein expression was induced with 0.5 mM IPTG for 3 h at 37°C. Cells were harvested by centrifugation and stored at −80°C. All subsequent procedures were performed at 4°C. Thawed bacteria were resuspended in 25 ml of buffer Y (50 mM Tris-HCl, pH 8.0, 500 mM NaCl, 10% glycerol) and supplemented with one tablet of protease inhibitor cocktail (Roche). The suspension was adjusted to 0.1 mg/ml lysozyme (Sigma) and incubated on ice for 30 min and the lysate was sonicated to reduce viscosity. Insoluble material was removed by centrifugation. The soluble extracts were loaded on a 2 ml of Strep-Tactin® Sepharose® resin (IBA, 2-1201-002) that had been equilibrated with buffer Y. The column was washed with 10 ml aliquots of buffer Y and then eluted five times with 1 ml aliquots of buffer Y containing 2.5 mM desthiobiotin (IBA). The first three fractions were concentrated dialyzed O/N in buffer Y at 4°C and concentrated using Amicon® Ultra-4 centrifugal filters (Merk).

Protein concentration determined using the Bio-Rad dye reagent with BSA as the standard. The final elution profiles were also monitored by SDS-PAGE (Fig S5).

### Co-immunoprecipitation assays

Purified *P. putida* histidine kinase (PP_HK, 10 µg) or its PAS domain (HK-PAS, 10 µg) were applied to 150 µl of Strep-Tactin® Sepharose® resin (IBA) and incubated in presence or absence of 25 µg of *P. putida* KaiC purified protein O/N at 4°C in buffer BB (50 mM Tris-HCl, pH 8.0, 200 mM NaCl, 0.01 % Triton X-100). As control for unspecific binding, KaiC proteins (25 µg) alone were added to 150 µl of Strep-Tactin® Sepharose® resin (IBA). Beads were washed 3 times with 150 µl buffer BB before resuspending in 100 µl Laemmli loading buffer. Each elution was then analyzed by SDS-PAGE and immunoblotting using anti-FLAG antibodies (Sigma, F3165) or anti-STREP antibodies (IBA, StrepMAB-Classic HRP conjugate) and following manufacturer protocol for immunodetection (https://www.iba-lifesciences.com/strepmab-classic-conjugate/2-1509-001).

### Graphics and Statistical analysis

Graphs were generated, and statistical analyses carried out, using the basic R graphical and statistic packages [30].

## Results

### *KaiC-like proteins are widespread in environmental* Pseudomonas

KaiB and KaiC homologs were searched in 1255 complete genomes from the *Pseudomonas* genus using Hidden Markov Models (hmm). While no KaiB-like protein was found, 222 KaiC homologs were retrieved and classified into three distinct phylogenetic groups (Fig. 1A). If we except *P. aeruginosa*, about 70% of *Pseudomonas* strains encode at least one KaiC homolog (Fig. 1B); among environmental species, only *Pseudomonas chlororaphis* and *Pseudomonas mendocina* overwhelmingly lack a KaiC homolog (SFile 1). Moreover, 17% of the strains encode more than one KaiC homolog, belonging in general to different phylogenetic groups; this suggests that different groups of KaiC homologs perform different functions.

The phylogenetic group I (dominant in *Pseudomonas fluorescens, Pseudomonas syringae*, and *P. protegens*) has a TY or SY pair at the site corresponding to the *Synechococcus* phosphorylated residues S431 and T432 (Fig. 1A, inclusions); the phylogenetic group III (dominant in *P. putida*) has an SS, sometimes a CS or SG, pair at the same position; the intermediate phylogenetic group II has no phosphorylatable residues at the corresponding position, but two threonines are conserved nearby which could be phosphorylated instead. Proteins from group II are almost always found in genomes which also contain a protein from group I (SFile 1), suggesting that group II KaiC homologs might crosstalk with group I KaiC homologs or be under the same selective pressures. Aside some sporadic gammaproteobacterial hits, the most closely related proteins outside of the *Pseudomonas* genus are found among Rhizobiales (alphaproteobacteria) for group I and III, and among Xanthomonadales (gammaproteobacteria) for group II. This hints at a possible role of KaiC homologs for growth on or around plants, since Rhizobiales and Xanthomonadales species are known plant symbionts and pathogens respectively.

KaiC homologs are usually found next to, and in the same predicted operon as, histidine kinases, hybrid histidine kinases, and/or response regulators. Typical genomic contexts of group I and group III KaiC homologs, exemplified in *P. protegens* CHA0 and *P. putida* KT2440 respectively, are shown in Figure 1C. To get a global overview of the neighboring genes, we identified the four proteins encoded closest to each *kaiC* gene using *ad hoc* hmm models and derived associations between a particular KaiC group and specific protein model in all genomes where only one KaiC homolog was found (Figure 1D, SFile 1). Group I KaiC homologs are usually found close to a histidine kinase, a single-domain response regulator, and a predicted EPS sortase; group III KaiC homologs are almost always found next to a histidine kinase, a single-domain response regulator, and a RNA-binding translational regulator of the CsrA-RsmA family; group II KaiC homologs are only associated with a histidine kinase. Altogether, these data suggest that KaiC homologs in *Pseudomonas* might often regulate or modulate gene expression through a histidine kinase, and a single-domain response regulator. Since response regulators associated with KaiC homologs do not directly bind DNA, they might act by interfering with the phosphorylation of actual DNA-binding regulators, functioning for instance as a phosphate sink.

### KaiC-like proteins are expressed during early stationary phase and become phosphorylated as stationary phase progresses

In order to characterize KaiC-like proteins in environmental *Pseudomonas* species, we chose to work on *P. protegens* CHA0 and *P. putida* KT2440, which encode a single KaiC-like protein from group I and III respectively. The genomic context of the genes encoding these KaiC homologs is shown in Figure 1C.

We first investigated whether KaiC-like proteins were expressed under standard culture conditions using a reporter construct, in which the promoter regions of each gene drive the transcription of a green fluorescent protein (GFP). While both reporters already showed significant transcriptional activity in exponential phase, the GFP signal peaked in early stationary phase in both species and remained stable thereafter (Fig. 2A and 2B). These results indicate that the maximal transcription rate of *kaiC* genes occurs in the beginning of stationary phase. To confirm that the cellular concentration of the KaiC-like protein is consistent with its transcription rate, we replaced the native *kaiC* sequence with a sequence coding for a FLAG-tagged KaiC protein. We evaluated the concentration of the protein at different time points during the culture using the immunoblot technique. The concentration of KaiC protein steadily increased from exponential to early stationary to advanced stationary phase for both *P. protegens* CHA0 and *P. putida* KT2440 (Fig. 2C). The KaiC protein therefore accumulates over stationary phase, which is consistent with the observed transcription pattern. These results suggest that KaiC proteins might help facing the challenges of life in stationary phase (under high cell density, nutrient limitation, or physiological stress).

**Figure 2:**
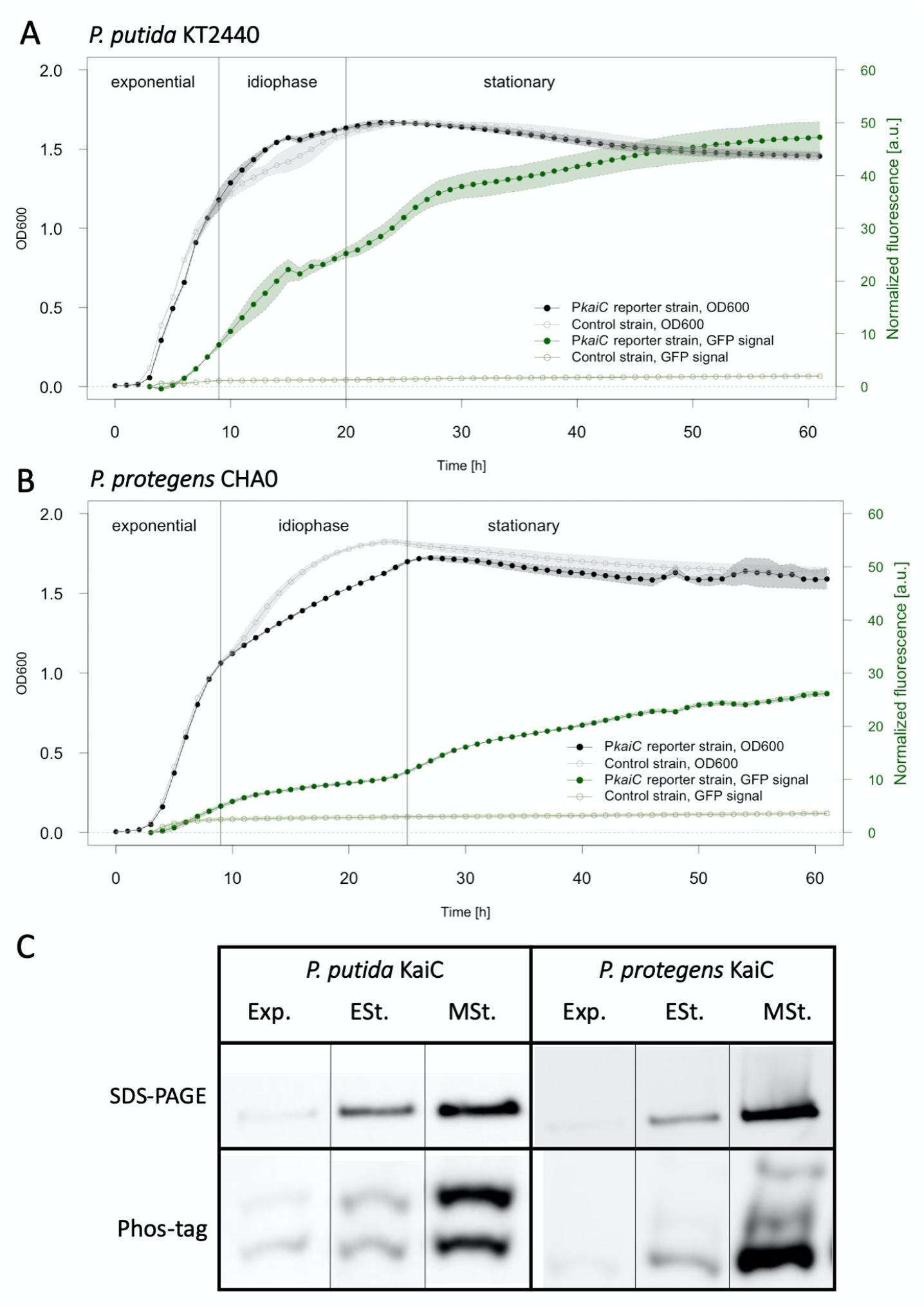
KaiC is highly expressed and phosphorylated in stationary phase. A. The *kaiC* gene is maximally transcribed at the transition from exponential phase to idiophase and at the transition from idiophase to stationary phase in *P. putida* KT2440. B. The *kaiC* gene is maximally transcribed at the transition from exponential phase to idiophase and at the transition from idiophase to stationary phase in *P. protegens* CHA0. C. The cellular concentration of FLAG-tagged KaiC, as detected by immunoblot after SDS-PAGE, is minimal in exponential phase (Exp.), but increases dramatically between early stationary phase (ESt., 12h) and mid-stationary phase (MSt., 24h) in *P. putida* KT2440 and *P. protegens* CHA0. Two phosphorylation states of the KaiC homolog could be observed in *P. putida* KT2440 from exponential phase on, while three phosphorylation states were detected in *P. protegens* CHA0 as of mid-stationary phase.

In the model cyanobacterium *S. elongatus*, the relative phosphorylation level of residues S431 and T432 varies depending on the circadian time. We wondered whether KaiC-like proteins from environmental *Pseudomonas*, in which phosphorylatable residues are at least partially conserved (Fig. 1A), could also be phosphorylated and whether phosphorylation would also vary over time. We used the Phos-tag™ system (Wako Chemicals) to detect different phosphorylation states of the KaiC-like proteins in a time course experiment. We observed two major phosphorylation states in the *P. putida* KT2440 homolog and three major phosphorylation states in the *P. protegens* CHA0 homolog (Fig. 2C). Moreover, in both species, the relative amounts of the alternative phosphorylation states varied over time during standard growth (Fig. 2C): hardly present in exponential phase, the one-time phosphorylated form of *P. putida* KT2440 KaiC-like protein is almost as abundant as the non-phosphorylated form in early and mid-stationary phase; by contrast, the *P. protegens* CHA0 KaiC homolog becomes two times phosphorylated only in mid-stationary phase. These results suggest that KaiC-like proteins in *Pseudomonas* are not immediately phosphorylated after being produced, but might become phosphorylated over time or in response to a specific trigger in early and mid-stationary phase.

### KaiC-like proteins are involved in environmental stress resistance

To investigate the function of KaiC-like proteins, we constructed a Δ*kaiC* deletion mutant of *P. protegens* CHA0 and *P. putida* KT2440. No difference in growth rate or yield was apparent during culture in rich or minimal media. In search of a phenotype, we characterized the transcriptome of cells grown in rich medium, hoping that differentially regulated genes would help us close-in on a potential function. We performed RNA-sequencing on cells collected in early, mid-, and late stationary phase for *P. putida* KT2440 and in late stationary phase only for *P. protegens* CHA0; the emphasis on stationary phase was chosen based on the expression and phosphorylation patterns observed in Figure 2C. Very limited overlap was observed between genes differentially regulated in the two species, suggesting that KaiC-like proteins of groups I and III are involved in different functions.

In *P. putida* KT2440, the distinctive expression pattern of the Δ*kaiC* mutant in early and mid-stationary phase was dominated by genes involved in stress resistance and catabolism of alternative carbon sources (SFile 2). Genes involved in glycine-betaine import and degradation on one hand and genes involved in the phenylacetic acid degradation pathway on the other hand were highly overexpressed in Δ*kaiC* in mid-stationary phase; a number of these genes were slightly, albeit significantly, overexpressed in early stationary phase. In late stationary phase, significant differences between the wild type and the Δ*kaiC* strain included *rpoS*, encoding the sigma factor helping cells adapt to stationary phase, the superoxide dismutase, involved in oxidative stress resistance, and genes involved in energy production through oxidation of fatty acids, which were both overexpressed. This indicates that the absence of KaiC might have different consequences depending on the age of the culture. Overall, these transcriptomic results suggest that the *P. putida* KT2440 KaiC homolog might help regulate the usage of alternative carbon and energy sources (glycine-betaine, phenylacetic acid, fatty acids, etc.) when nutrients become limiting.

In late stationary phase, the Δ*kaiC* mutant strain of *P. protegens* CHA0 showed repressed import of sulfur compounds and repressed catabolism of alternative carbon sources, like amino-acids, aromatic compounds, or polysaccharides (SFile 2). Several transcription factors were also dysregulated. This pattern fits with an involvement of the *P. protegens* CHA0 KaiC homolog in adaptation to stationary phase.

### *In* P. putida *KT2440, KaiC protects against osmotic and oxidative stresses*

Based on the transcription pattern of the *kaiC* gene and the global pattern of the RNA-sequencing data, we hypothesized that KaiC might be involved in resistance to physiological stresses. We therefore tested survival after 72h of growth, growth at high (37°C) or low (18°C) temperatures, growth under red or blue light, growth under oxidative stress (increasing concentrations of H_2_O_2_), and growth under osmotic stress (increasing concentrations of NaCl) on *P. putida* KT2240 and *P. protegens* CHA0. No difference between the wild-type and the Δ*kaiC* mutant was observed in any of the tested conditions for *P. protegens* CHA0 (data not shown). Temperature, light, and long-term starvation did not have a differential impact on the wild type and Δ*kaiC* mutant in *P. putida* KT2440 (data not shown). However, the *P. putida* KT2440 Δ*kaiC* mutant presented a higher sensitivity to H_2_O_2_ both in rich and in minimal media (Fig. 3A). In addition, a different growth pattern was apparent at relatively high NaCl concentrations (1 and 2 M) between the wild type and the Δ*kaiC* mutant with the Δ*kaiC* mutant starting its growth later (longer lag phase) (Fig. 3B, left). This effect was strongly attenuated upon addition of glycine-betaine (10 µg/ml) (Fig. 3B, right). This indicates that, in the absence of KaiC, *P. putida* KT2440 adapts less efficiently to high osmolarity growth conditions. These results are consistent with the untimely activation of glycine-betaine catabolism in early to mid-stationary phase observed in the RNA-sequencing data, since this small molecule is one of the main osmoprotectants in bacteria.

**Figure 3:**
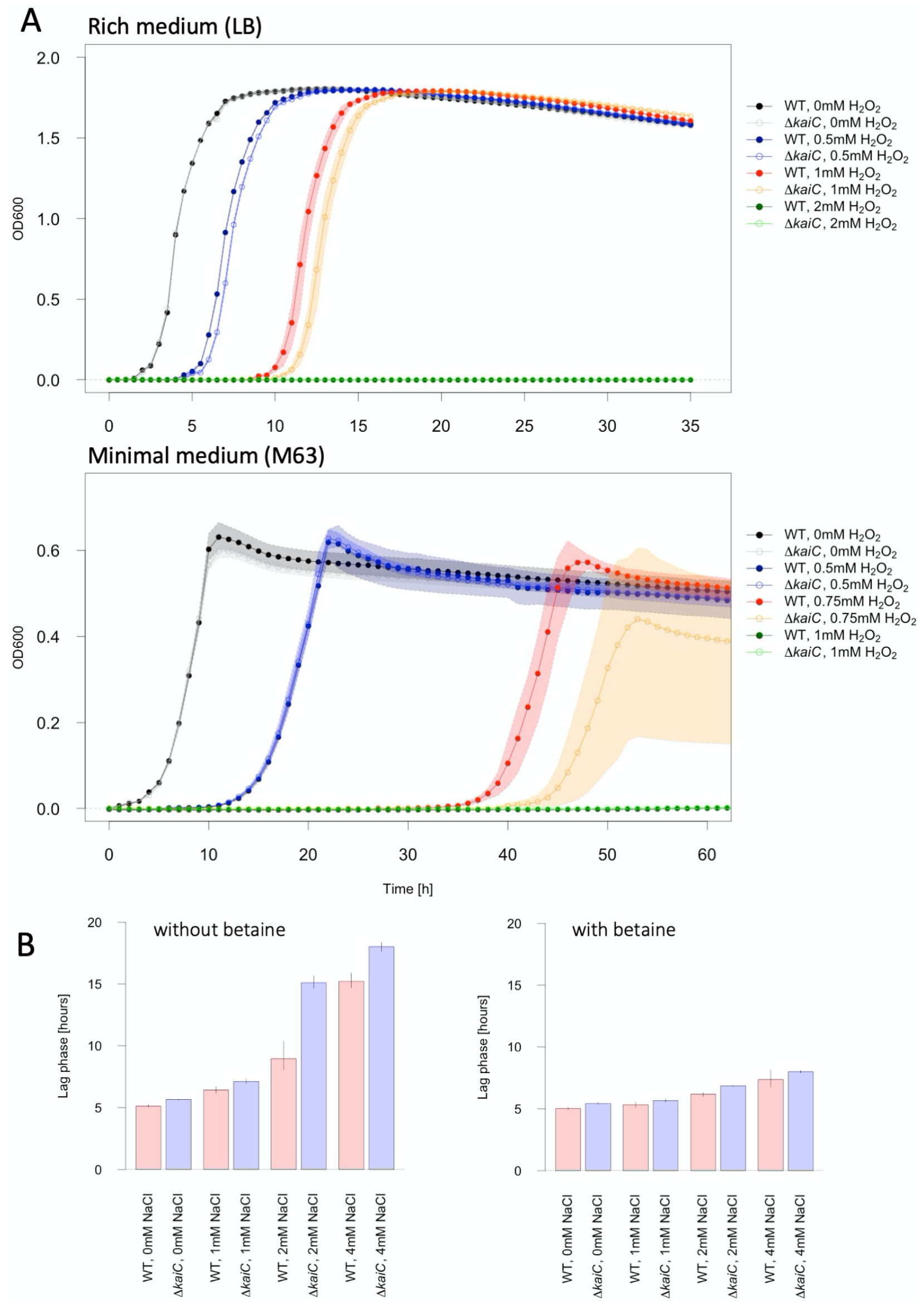
In *P. putida* KT2440, the KaiC homolog is required for maximal protection against oxidative and osmotic stresses. A. In *P. putida* KT2440, the Δ*kaiC* mutant strain is less resistant to oxidative stress (increasing H_2_O_2_ concentrations) than the wild type in rich (upper part) and minimal medium (lower part). B. In *P. putida* KT2440, the Δ*kaiC* mutant strain has a longer lag-phase than the wild type strain under high NaCl concentrations in minimal medium (right); this phenotype is mostly suppressed if 10 µg/ml glycine-betaine are added to the medium (left).

### KaiC interaction network

In Cyanobacteria, KaiC is a protein interaction hub: KaiA and KaiB bind KaiC to tune and time its auto-kinase/auto-phosphatase activity, while kinase SasA and phosphatase CikA are activated or deactivated upon KaiC-binding. We asked whether the proteins encoded by genes most often found in the neighborhood of KaiC homologs in *Pseudomonas* species physically interacted with it. To this purpose, we tested the *P. putida* KT2440 and *P. protegens* CHA0 KaiC homologs and their neighbors (the hybrid histidine kinase and response regulator in *P. putida* KT2440, the histidine kinase in *P. protegens* CHA0) in a heterologous two-hybrid system (Fig. 4ABC, SFig. 1AB and 2AB). The system is based on the expression in *E. coli* of two proteins of interest that are cloned and in frame with either T25 or T18 split adenylyl cyclase domains [34]. Domains proximity, due to protein-protein interactions, restores the enzyme activity and results in red colonies when cells were grown in Congo Red medium or in a significant increase of β-galactosidase activity compared to controls. We found that KaiC strongly interacted with itself in both *Pseudomonas* species, which is consistent with its predicted hexameric quaternary structure. Both KaiC homologs interacted with their respective histidine kinase, strongly in *P. protegens* and significantly but much more weakly in *P. putida*. In *P. putida*, interactions between KaiC and the response regulator on the one hand and between the histidine kinase and the response regulator on the other hand were also positive. Both histidine kinases interact with themselves, likely as dimers. We confirmed *in vitro* the interaction between *P. putida* KaiC and the histidine kinase (HK) by purifying the proteins separately from *E. coli* (SFig. 5) and performing co-immunoprecipitation assays. Fig. 4D shows that His-tagged KaiC specifically co-immunoprecipitated with Strep-tagged HK. This interaction most likely occurs through the N-terminal PAS (Per-Arnt-Sim) domain of the histidine kinase since this domain is sufficient for interaction. Overall, these results suggest that KaiC homologs in *Pseudomonas* species act as a multimer (most likely a hexamer) through interacting partners involved in phosphotransfer signaling networks (Fig. 4E).

**Figure 4:**
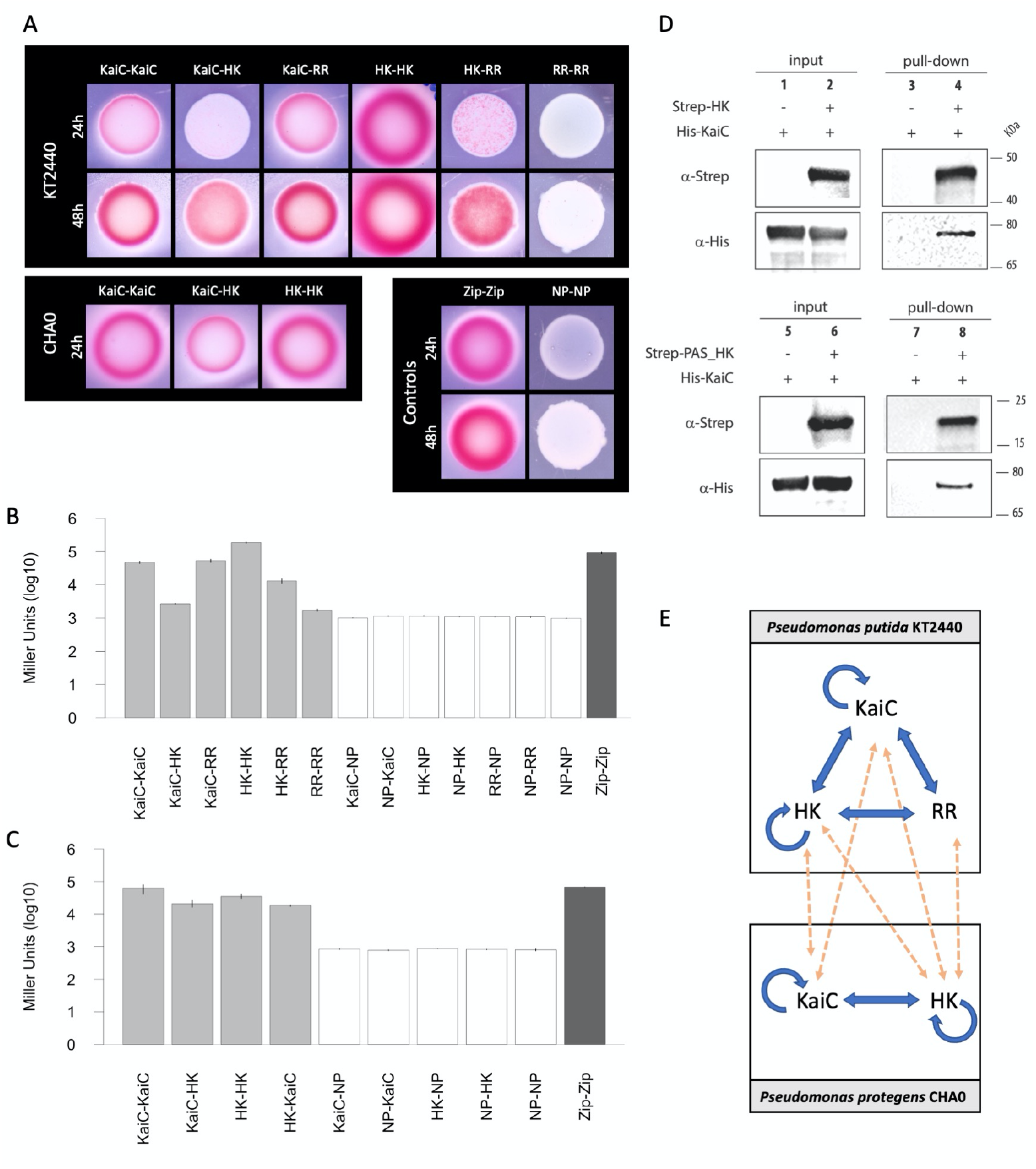
KaiC interactions network in *P. putida* KT2440 and *P. protegens* CHA0. A. Two-hybrid spot assay for KaiC interactions in KT2440 and CHA0. The intensity of the pink color is roughly proportional to the interaction strength of the two proteins tested. HK: histidine kinase; RR: response regulator; NP: no protein; Zip: leucine zipper, positive control. B. β-galactosidase assay quantifying the interactions between KaiC and KaiC-associated proteins in *P. putida* KT2440 24h after spotting. C. β-galactosidase assay quantifying the interactions between KaiC and KaiC-associated proteins in *P. protegens* CHA0 24h after spotting. D. Immunoblot of before (input) and after co-immunoprecipitation (Co-IP), showing that KaiC and its cognate histidine kinase (HK) interact *in vitro* and that the PAS-domain of the protein is sufficient for interaction. E. Model of KaiC interactions network in KT2440 and CHA0; yellow dotted arrows show positive cross-interactions between the two systems. E. Model of KaiC interactions network in KT2440 and CHA0; yellow dotted arrows show positive cross-interactions between the two systems.

Interestingly, cross-interactions between the homologs of *P. putida* KT2440 and *P. protegens* CHA0 were also observed for the KaiC proteins and the histidine kinases, as well as between the response regulator of *P. putida* KT2440 and the histidine kinase of *P. protegens* CHA0 (SFig. 3 and 4). This suggests that when more than one KaiC protein are present in a single genome these proteins could cross-interact and cross-talk with one another (Fig. 4E).

## Discussion

How the sophisticated circadian clock found in Cyanobacteria evolved is a long standing question. Homologues of the KaiC protein, the main driver of the cyanobacterial clock, can be found, beyond Cyanobacteria, in many archaeal and bacterial genomes. Here, we contribute to shedding light on the diversity and function of KaiC-like proteins in non-photosynthetic bacteria. We focus on environmental *Pseudomonas* species, which are often studied for their pollutant-degrading or plant growth promoting properties.

Our results show that KaiC homologs belonging to three main phylogenetic groups are widespread among environmental *Pseudomonas* species. Almost all strains in the “*putida”* and in the “*syringae”* group and more than half of the strains in the “*fluorescens”* and “*stutzeri”* groups encode at least one KaiC homolog, with 20% of KaiC-encoding species encoding more than one homolog (Fig. 1AB, SFile 1). KaiC-like proteins most likely provide a substantial selective advantage to these environmental species. However, co-conserved genes do not give any indication about the function they might perform, as no physiologically characterized enzymes are systematically found in the neighborhood. Nevertheless, KaiC homologs are virtually always associated with a histidine kinase and sometimes single-domains response regulators, which suggests that they might participate in regulatory networks. We chose to characterize the expression, function, and physical interaction network of the homologs present in *P. putida* KT2440 and *P. protegens* CHA0 as representatives of the two most abundant groups of KaiC-like proteins.

We found that *P. putida* KT2440 and *P. protegens* CHA0 homologs are both most abundantly expressed in early stationary phase and that their phosphorylation state changes as stationary phase progresses (Fig. 2). This suggests that their function is linked to conditions in which nutrients become limiting and physiological stress increases. Based on the cyanobacterial KaiC model, changes in the phosphorylation state of the protein during stationary phase could induce alternative conformations, leading to different physical interactions and functional outputs. In this perspective, KaiC-like proteins in *Pseudomonas* species could function as starvation or stress “timers”, fine-tuning stress-resistance networks in a time-dependent manner. Consistent with this hypothesis, the transcription profile of *P. putida* Δ*kaiC* markedly changes between early or mid-stationary phase on one hand and late stationary phase on the other hand. Protein populations dominated by the non-phosphorylated form (early or mid-stationary phase) could therefore lead to different phenotypes as compared to protein populations dominated by phosphorylated forms (late stationary phase). This could be compared with the “hourglass” function proposed for the *Rhodopseudomonas palustris* KaiC homolog [9]; however, instead of timing transcription over a 24h time period, it would measure the duration of stress periods and help with adaptations over longer time scales.

The co-conservation (Fig. 1CD) and physical interaction (Fig. 4) of KaiC homologs with a histidine kinase (*P. putida* and *P. protegens* homologs) and a single-domain response regulator (*P. putida* homolog), suggest that KaiC proteins might play a role in phospho-transfer networks and indirectly regulate transcription. The KaiC-associated histidine kinases might directly phosphorylate (or dephosphorylate) unidentified DNA-binding transcriptional regulators in a KaiC-dependent manner, by being spatially sequestered or activated/deactivated when bound to KaiC. Likewise, single domain response regulators can be used as phosphate sink or source within phosphotransfer networks and indirectly influence the phosphorylation state of multiple players, including transcriptional regulators [35]. This would be consistent with the relatively small transcriptional effects, in terms of number of affected genes and fold-change, observed in the Δ*kaiC* mutants in both strains. The KaiC homolog in *P. putida*, for instance, seems to change the temporal dynamics of betaine-utilization gene transcription, fostering earlier transcription and more limited accumulation of the transcript in early stationary phase. However, the effect-size remains small in RT-qPCR and reporter assays.

The transcriptional profile of the Δ*kaiC* mutant in *P. putida* and P. protegens reveals significant expression changes in genes involved in stress resistance (osmotic stress, oxidative stress, sulfur starvation) and alternative carbon source catabolism (short chain fatty acids, aromatic compounds). Such functions could be important to withstand harsh environmental conditions (stress and starvation). Interestingly, glycine-betaine metabolism, which appears impacted in all three sets of RNA-sequencing data in *P. putida*, implies a trade-off between nutrition and stress resistance since glycine-betaine is, in bacteria, both a carbon source and an osmoprotectant [36]. The right balance between glycine-betaine degradation or its usage as an osmoprotectant likely changes as stationary phase progresses and carbon/energy sources become scarce. Our data suggest that the KaiC homolog in *P. putida* could prevent glycine-betaine degradation in early stationary phase (promoting its osmoprotective function), but not in late stationary phase when energy and carbon become crucial. Likewise, sulfur starvation responses might need to be activated at a specific time during stationary phase in order to replenish the pools of sulfur-containing amino-acids. Overall, this supports the hypothesis that the KaiC homolog might be involved in fine-tuning or balancing different usages of common substrates.

Two crucial questions remain open in our model: 1) What are the intermediates between the KaiC-associated histidine kinase (and response regulator) and the observed transcriptional effects? and 2) What is the input given or received by KaiC? Does KaiC measure time, for instance through a condition-stable autophosphorylation kinetics, or does it sense something in the cell, e.g. ATP levels, that triggers or stimulates its autophosphorylation? These two questions should be the priority for future research.

## Acknowledgments

*Pseudomonas protegens* CHA0, *E. coli* DH5α-λpir as well as pEMG and pSW-2 plasmids were kindly provided by Christoph Keel lab from the University of Lausanne. *Pseudomonas putida* KT2440 was kindly provided by Dr. Ricardo Machado from the University of Neuchâtel. This project is supported by the Swiss National Science Foundation grant PZ00P3_180142.

## Supplementary figures

**SFigure 1:**
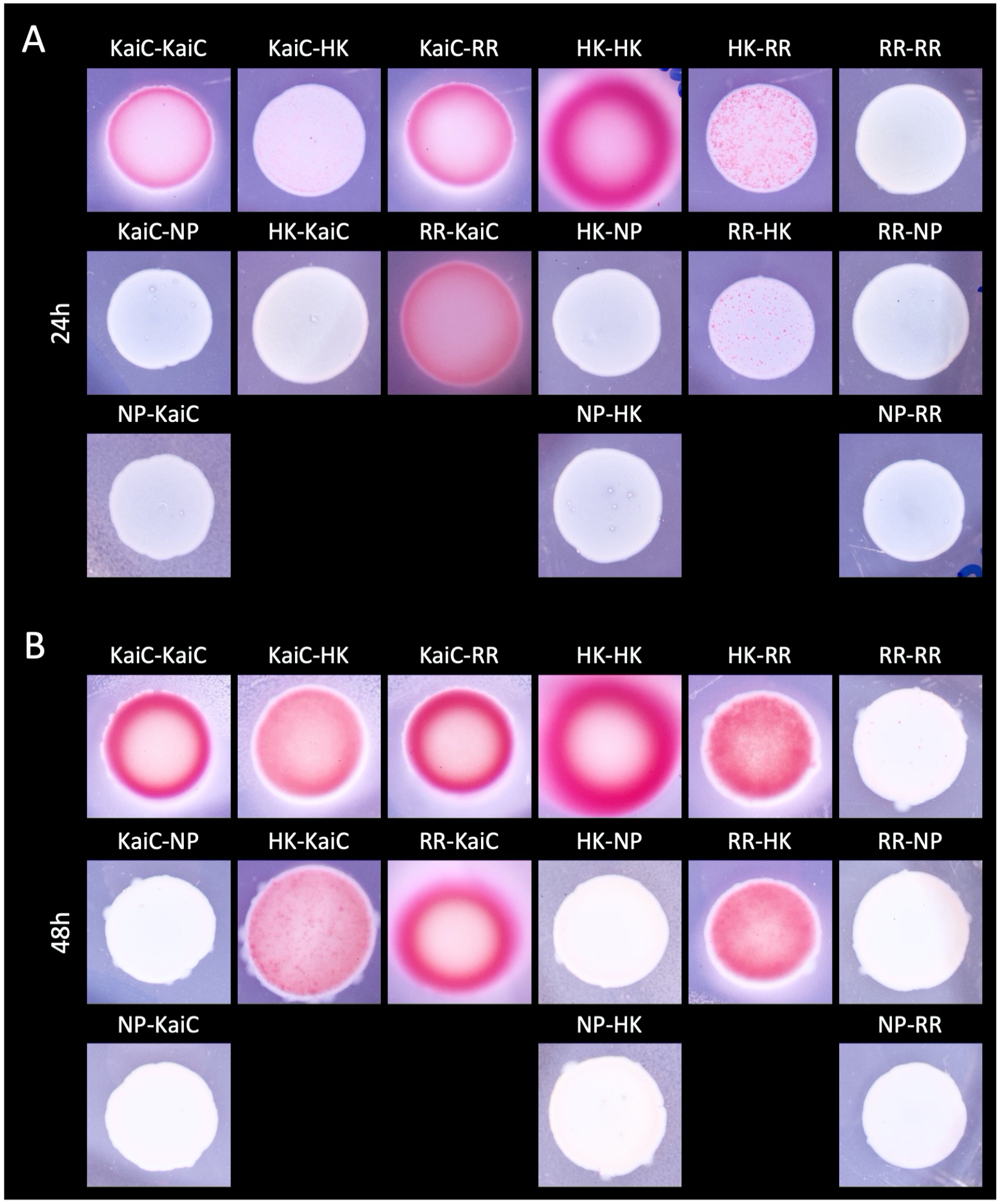
KaiC interactions network in *P. putida* KT2440 after 24h or 48h. A. Two-hybrid spot assay for KaiC interactions in KT2440 after 24h. B. Two-hybrid spot assay for KaiC interactions in *P. putida* KT2440 after 49h. The intensity of the pink color is roughly proportional to the interaction strength of the two proteins tested. HK: histidine kinase; RR: response regulator; NP: no protein; Zip: leucine zipper, positive control.

**SFigure 2:**
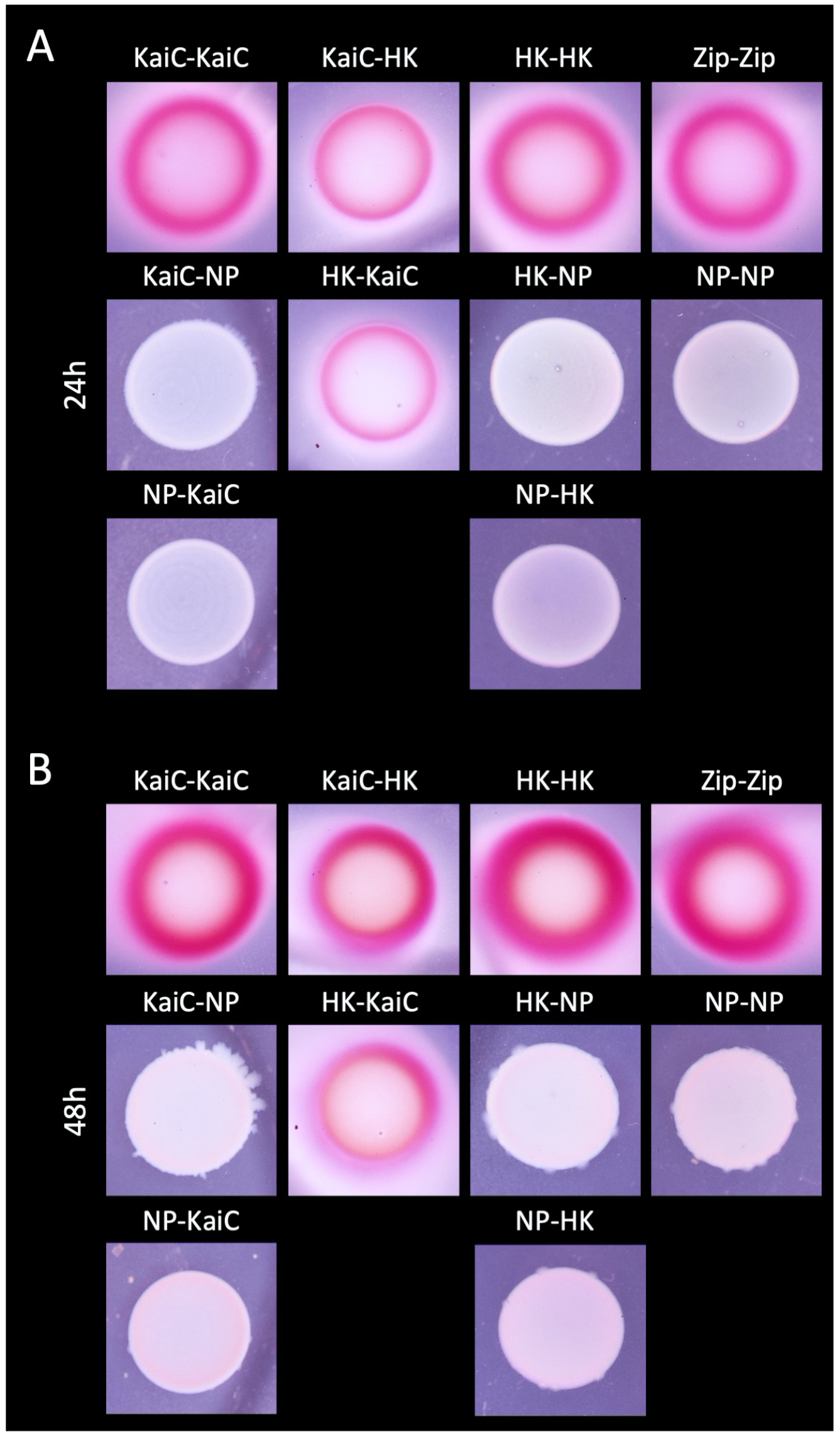
KaiC interactions network in *P. protegens* CHA0 after 24h or 48h. A. Two-hybrid spot assay for KaiC interactions in *P. protegens* CHA0 after 24h. B. Two-hybrid spot assay for KaiC interactions in *P. protegens* CHA0 after 49h. The intensity of the pink color is roughly proportional to the interaction strength of the two proteins tested. HK: histidine kinase; RR: response regulator; NP: no protein; Zip: leucine zipper, positive control.

**SFigure 3:**
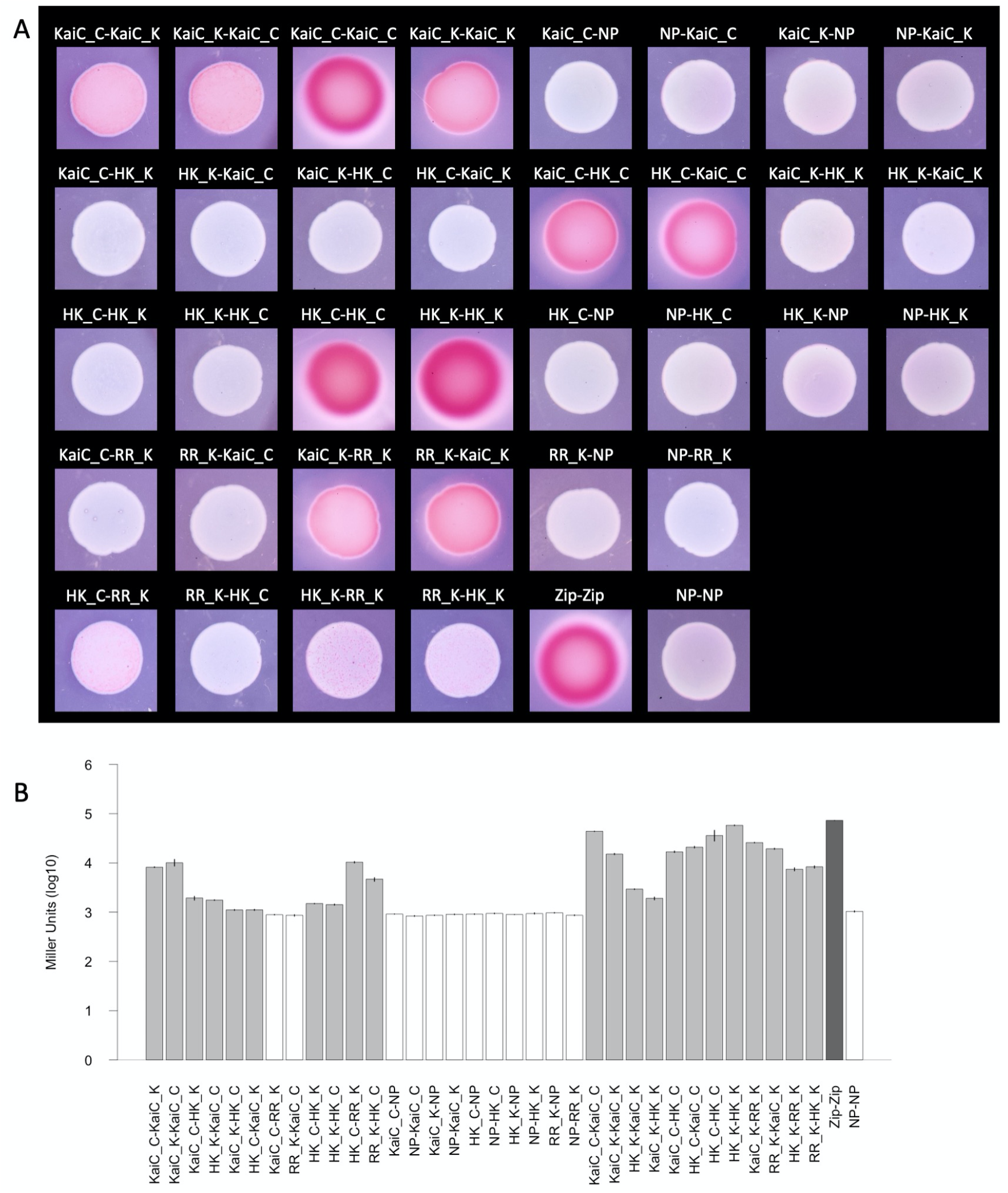
Cross-interactions in the KaiC network between *P. putida* KT2440 and *P. protegens* CHA0 after 24h. A. Two-hybrid spot assay for KaiC interactions in KT2440 after 24h. The intensity of the pink color is roughly proportional to the interaction strength of the two proteins tested. B. β-galactosidase assay quantifying the interactions between KaiC and KaiC-associated proteins from *P. putida* KT2440 and *P. protegens* CHA0 24h after spotting. Suffix “_K”: protein from *P. putida* KT2440; suffix “_C”: protein from *P. protegens* CHA0; HK: histidine kinase; RR: response regulator; NP: no protein; Zip: leucine zipper, positive control. B. β-galactosidase assay quantifying the interactions between KaiC and KaiC-associated proteins in *P. protegens* CHA0 after 24h.

**SFigure 4:**
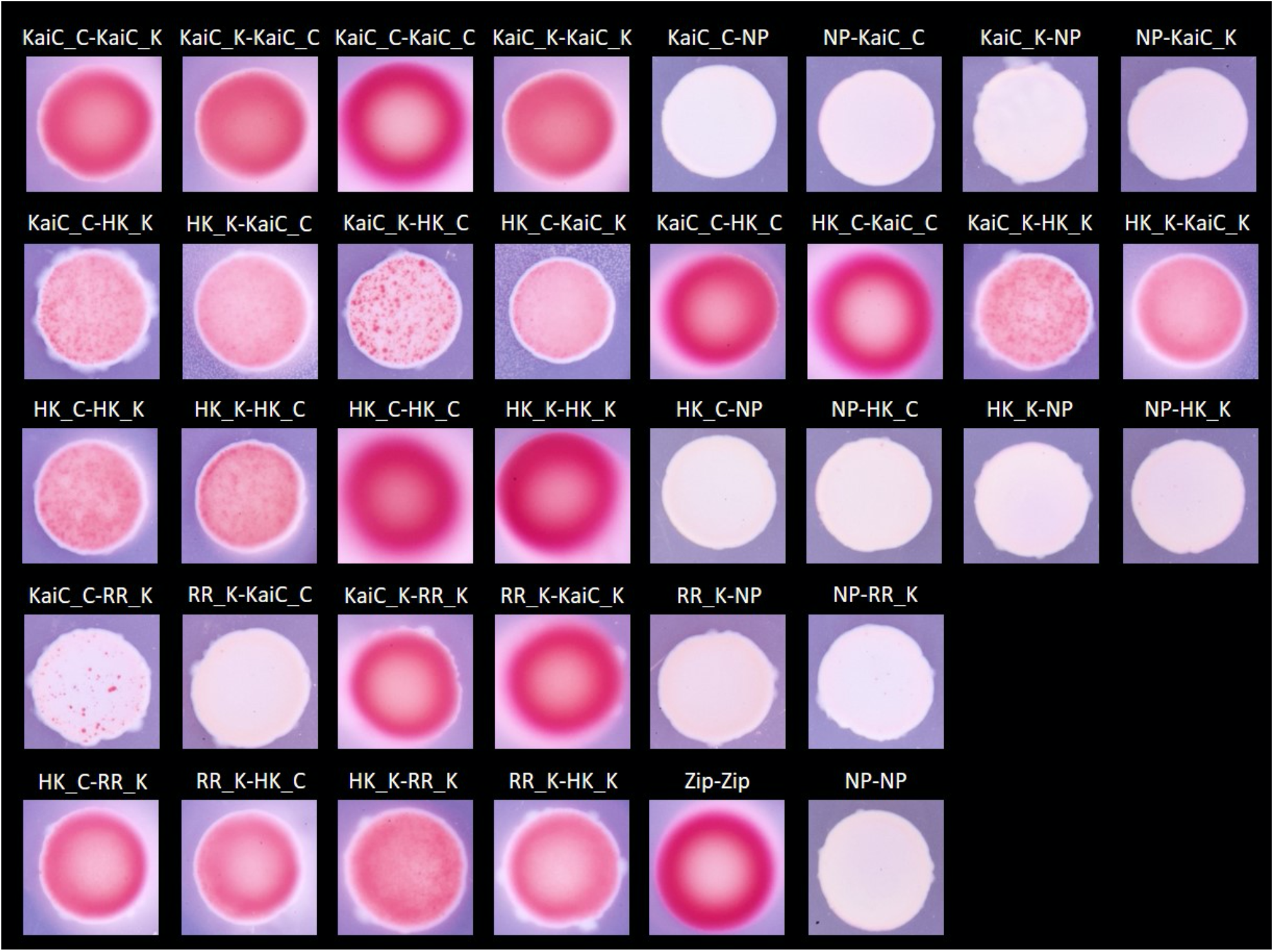
Cross-interactions in the KaiC network between *P. putida* KT2440 and *P. protegens* CHA0 after 48h. Two-hybrid spot assay for cross-interactions between proteins from *P. putida* KT2440 and *P. protegens* CHA0. The intensity of the pink color is roughly proportional to the interaction strength of the two proteins tested. Suffix “_K”: protein from *P. putida* KT2440; suffix “_C”: protein from *P. protegens* CHA0; HK: histidine kinase; RR: response regulator; NP: no protein; Zip: leucine zipper, positive control.

**SFigure 5:**
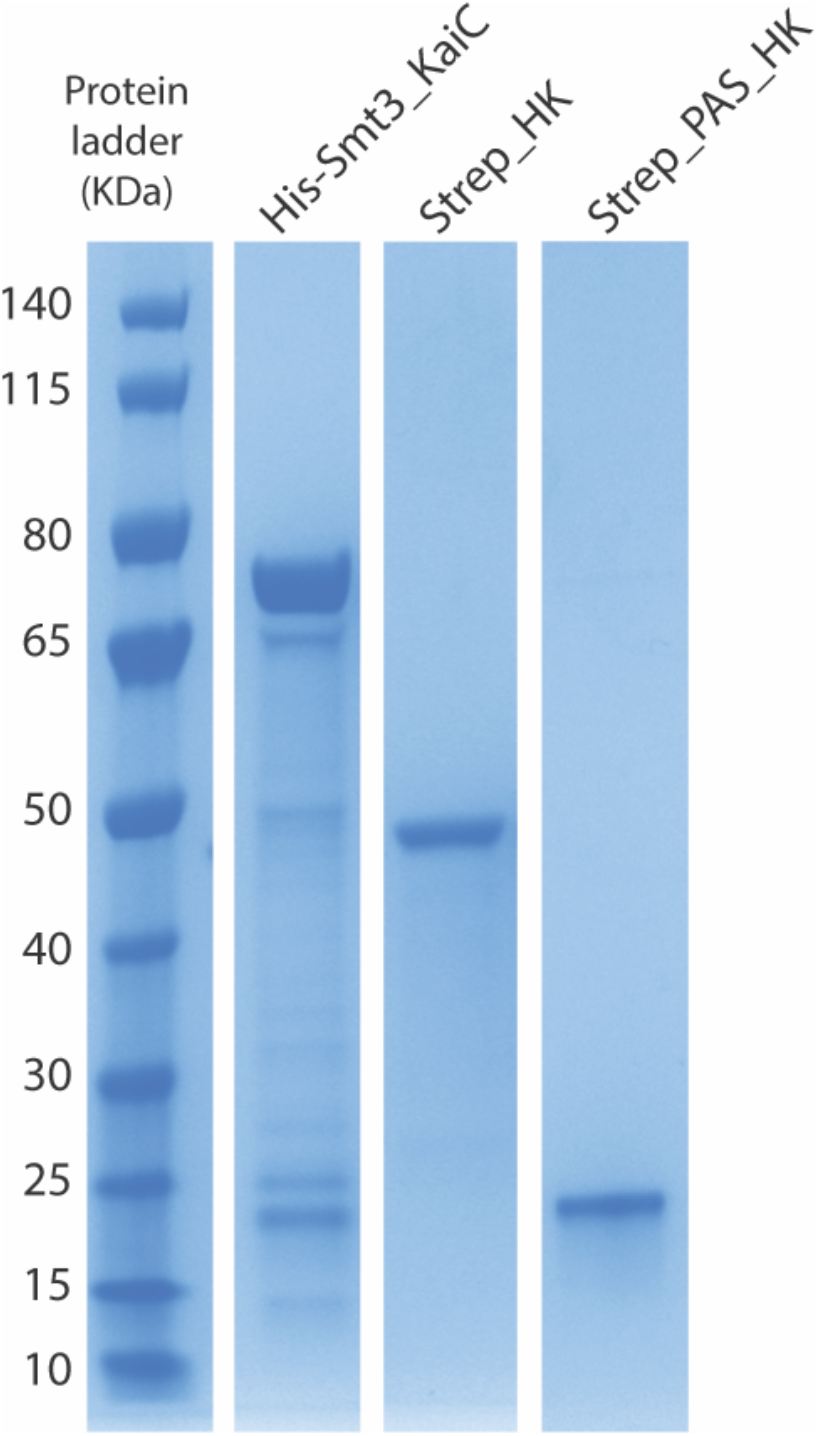
Purification of *P. putida* KT2440 KaiC (PP_3834), Histidine kinase (PP_3835) and its PAS domain (PP_3835, amino acid 1-138). Aliquots (2.5 µg) of the nickel agarose preparations of *P. putida* KT2440 KaiC (His-Smt3_KaiC), or of the strep sepharose resin of *P. putida* histidine kinase (Step_HK) and its PAS domain (Strep_PAS_HK) were analyzed by SDS-PAGE. Coomassie Blue-stained gel is shown. The positions and sizes (in kDa) of marker proteins are indicated on the left.

## Supplementary tables

**Table 1:**
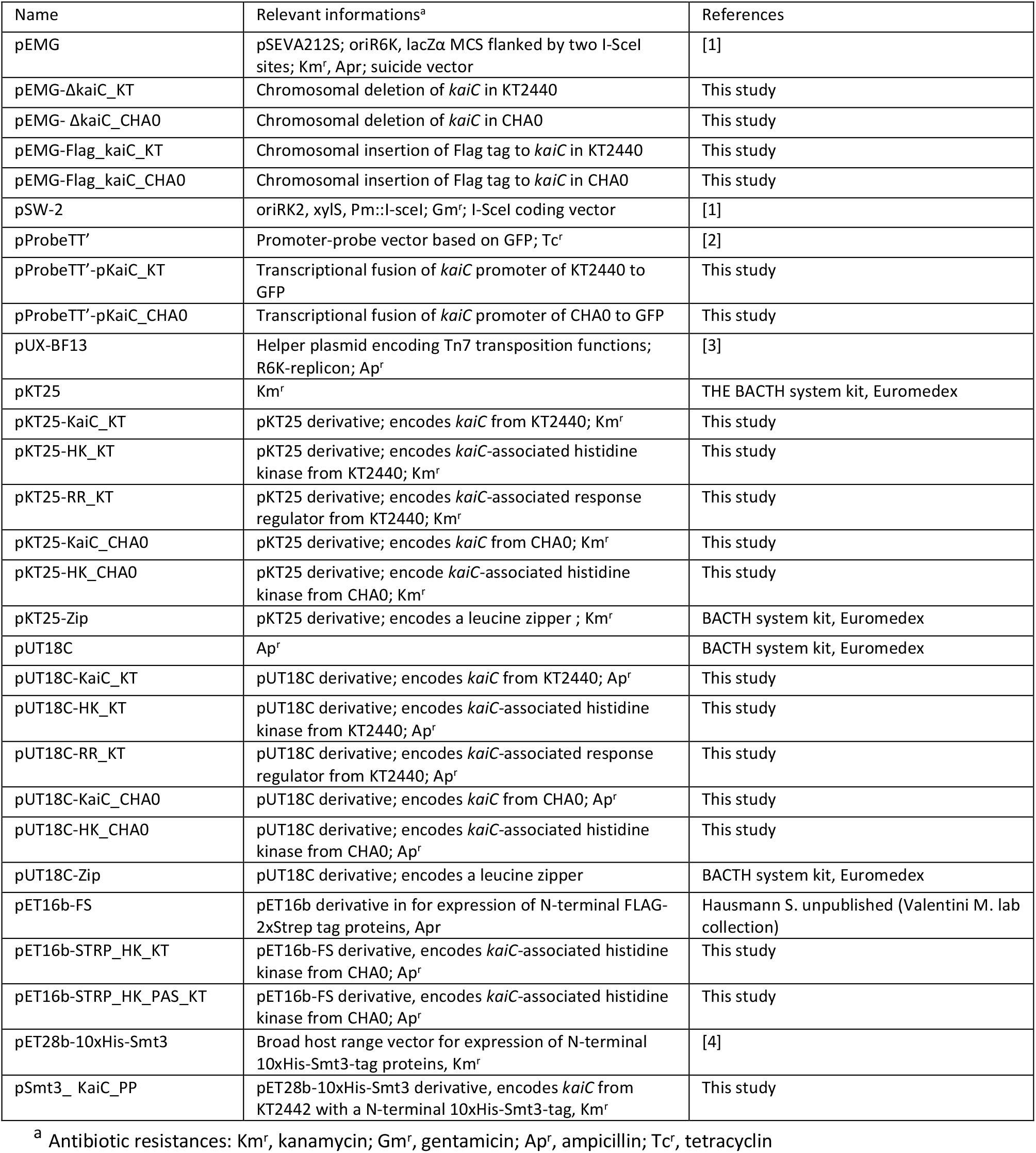
Plasmids used in this study

**Table 2:**
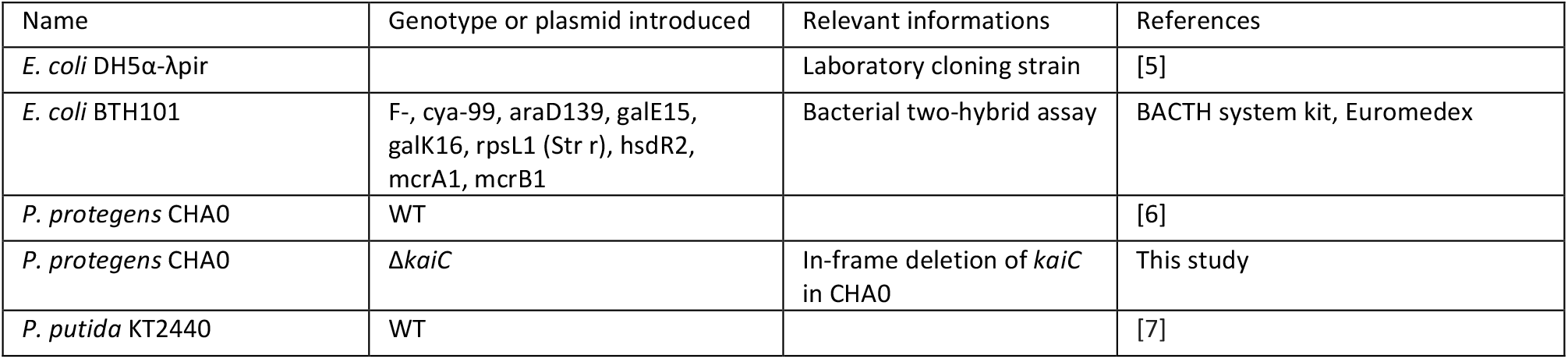
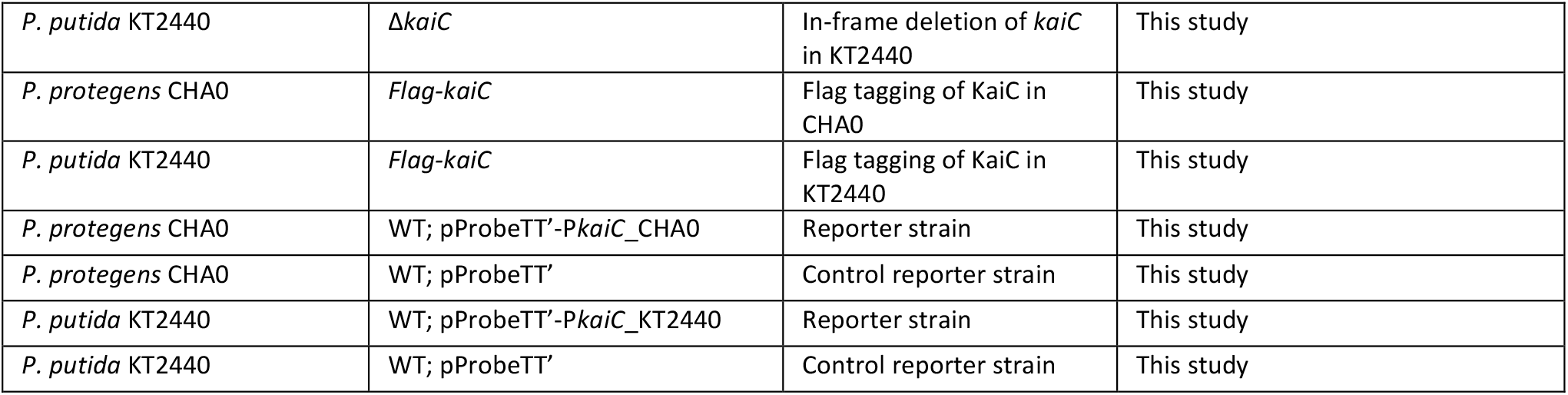
Strains used in this study

**Table 3:**
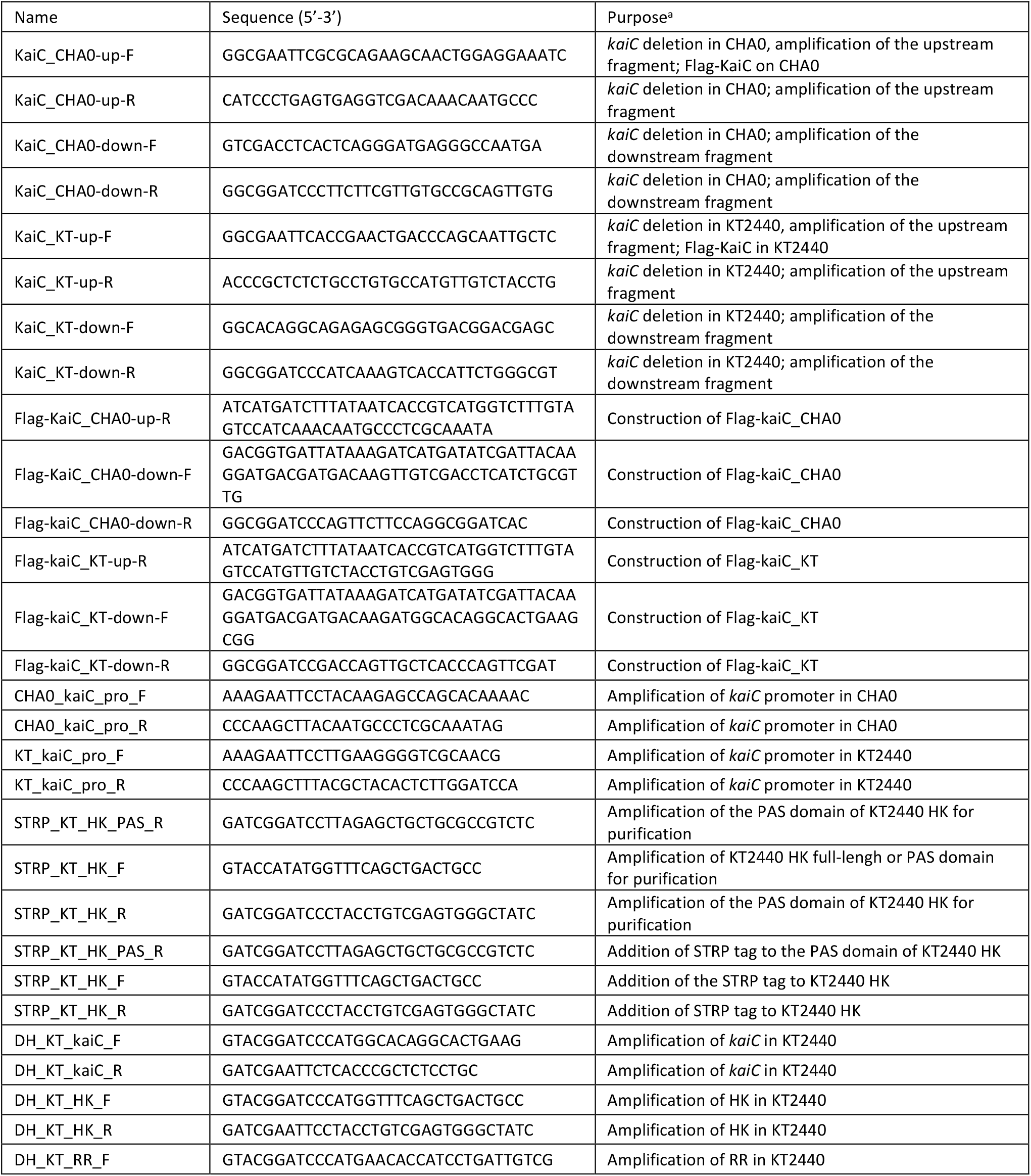
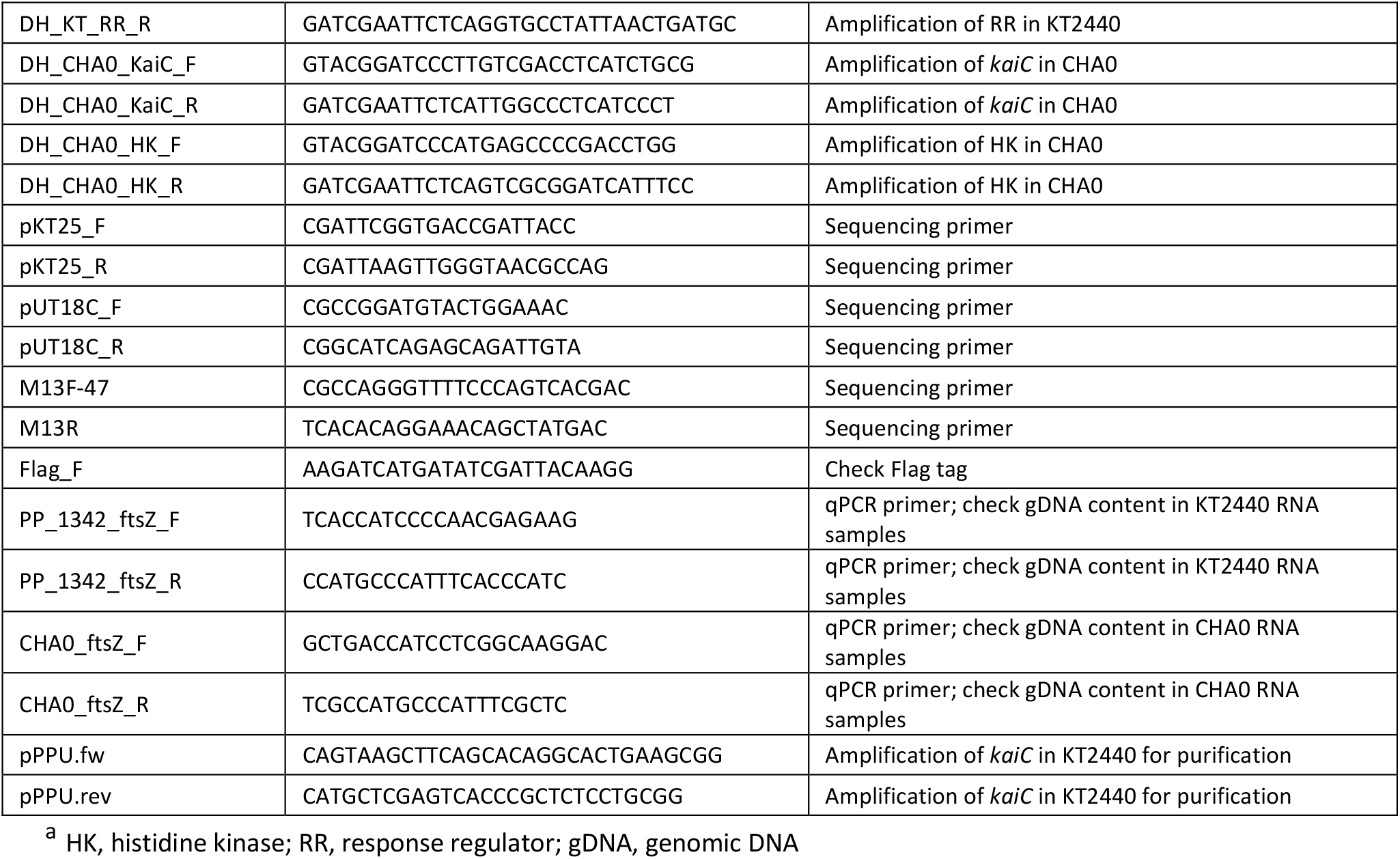
Primers used in this study

## References

1. Johnson CH, Stewart PL, Egli M: The Cyanobacterial Circadian System: From Biophysics to Bioevolution. Annu Rev Biophys 2011, 40:143–167.

2. Cohen SE, Golden SS: Circadian Rhythms in Cyanobacteria. Microbiol Mol Biol Rev 2015, 79:373–385.

3. Axmann IM, Hertel S, Wiegard A, Dörrich AK, Wilde A: Diversity of KaiC-based timing systems in marine Cyanobacteria. Mar Genomics 2014, 14:3–16.

4. Markson JS, Piechura JR, Puszynska AM, O’Shea EK: Circadian Control of Global Gene Expression by the Cyanobacterial Master Regulator RpaA. Cell 2013, 155:1396–1408.

5. Vijayan V, Zuzow R, O’Shea EK: Oscillations in supercoiling drive circadian gene expression in cyanobacteria. Proc Natl Acad Sci 2009, 106:22564–22568.

6. Dvornyk V, Vinogradova O, Nevo E: Origin and evolution of circadian clock genes in prokaryotes. Proc Natl Acad Sci 2003, 100:2495–2500.

7. Loza-Correa M, Gomez-Valero L, Buchrieser C: Circadian Clock Proteins in Prokaryotes: Hidden Rhythms? Front Microbiol 2010, 1.

8. Schmelling NM, Lehmann R, Chaudhury P, Beck C, Albers S-V, Axmann IM, Wiegard A: Minimal tool set for a prokaryotic circadian clock. BMC Evol Biol 2017, 17:169.

9. Ma P, Mori T, Zhao C, Thiel T, Johnson CH: Evolution of KaiC-Dependent Timekeepers: A Proto-circadian Timing Mechanism Confers Adaptive Fitness in the Purple Bacterium Rhodopseudomonas palustris. PLOS Genet 2016, 12:e1005922.

10. Maniscalco M, Nannen J, Sodi V, Silver G, Lowrey PL, Bidle KA: Light-dependent expression of four cryptic archaeal circadian gene homologs. Front Microbiol 2014, 5.

11. Iguchi H, Yoshida Y, Fujisawa K, Taga H, Yurimoto H, Oyama T, Sakai Y: KaiC family proteins integratively control temperature-dependent UV resistance in Methylobacterium extorquens AM1: KaiC as integrative controller of TDR. Environ Microbiol Rep 2018, 10:634–643.

12. Stutz EW: Naturally Occurring Fluorescent Pseudomonads Involved in Suppression of Black Root Rot of Tobacco. Phytopathology 1986, 76:181.

13. Kupferschmied P, Maurhofer M, Keel C: Promise for plant pest control: root-associated pseudomonads with insecticidal activities. Front Plant Sci 2013, 4.

14. Nikel PI, de Lorenzo V: Pseudomonas putida as a functional chassis for industrial biocatalysis: From native biochemistry to trans-metabolism. Metab Eng 2018, 50:142–155.

15. Wu X, Monchy S, Taghavi S, Zhu W, Ramos J, van der Lelie D: Comparative genomics and functional analysis of niche-specific adaptation in Pseudomonas putida. FEMS Microbiol Rev 2011, 35:299–323.

16. Regenhardt D, Heuer H, Heim S, Fernandez DU, Strompl C, Moore ERB, Timmis KN: Pedigree and taxonomic credentials of Pseudomonas putida strain KT2440. Environ Microbiol 2002, 4:912–915.

17. Wu L, McGrane RS, Beattie GA: Light Regulation of Swarming Motility in Pseudomonas syringae Integrates Signaling Pathways Mediated by a Bacteriophytochrome and a LOV Protein. mBio 2013, 4.

18. Sumi S, Mutaguchi N, Ebuchi T, Tsuchida H, Yamamoto T, Suzuki M, Natsuka C, Shiratori-Takano H, Shintani M, Nojiri H, et al.: Light Response of Pseudomonas putida KT2440 Mediated by Class II LitR, a Photosensor Homolog. J Bacteriol 2020, 202.

19. Mukherjee S, Jemielita M, Stergioula V, Tikhonov M, Bassler BL: Photosensing and quorum sensing are integrated to control Pseudomonas aeruginosa collective behaviors. PLOS Biol 2019, 17:e3000579.

20. Martínez-García E, de Lorenzo V: Engineering multiple genomic deletions in Gram-negative bacteria: analysis of the multi-resistant antibiotic profile of Pseudomonas putida KT2440: Tools for editing Gram-negative genomes. Environ Microbiol 2011, 13:2702–2716.

21. Miller WG, Leveau JHJ, Lindow SE: Improved gfp and inaZ Broad-Host-Range Promoter-Probe Vectors. Mol Plant-Microbe Interactions® 2000, 13:1243–1250.

22. Edgar RC: MUSCLE: multiple sequence alignment with high accuracy and high throughput. Nucleic Acids Res 2004, 32:1792–1797.

23. Price MN, Dehal PS, Arkin AP: FastTree 2 – Approximately Maximum-Likelihood Trees for Large Alignments. PLoS ONE 2010, 5:e9490.

24. Garrido-Sanz D, Meier-Kolthoff JP, Göker M, Martín M, Rivilla R, Redondo-Nieto M: Genomic and Genetic Diversity within the Pseudomonas fluorescens Complex. PLOS ONE 2016, 11:e0150183.

25. Thomsen MCF, Nielsen M: Seq2Logo: a method for construction and visualization of amino acid binding motifs and sequence profiles including sequence weighting, pseudo counts and two-sided representation of amino acid enrichment and depletion. Nucleic Acids Res 2012, 40:W281–W287.

26. Fu L, Niu B, Zhu Z, Wu S, Li W: CD-HIT: accelerated for clustering the next-generation sequencing data. Bioinformatics 2012, 28:3150–3152.

27. Chen S, Zhou Y, Chen Y, Gu J: fastp: an ultra-fast all-in-one FASTQ preprocessor. Bioinformatics 2018, 34:i884–i890.

28. Langmead B, Salzberg SL: Fast gapped-read alignment with Bowtie 2. Nat Methods 2012, 9:357–359.

29. Liao Y, Smyth GK, Shi W: featureCounts: an efficient general purpose program for assigning sequence reads to genomic features. Bioinformatics 2014, 30:923–930.

30. R Core Team: R: A Language and Environment for Statistical Computing. R Foundation for Statistical Computing; 2021.

31. Love MI, Huber W, Anders S: Moderated estimation of fold change and dispersion for RNA-seq data with DESeq2. Genome Biol 2014, 15:550.

32. Schneider CA, Rasband WS, Eliceiri KW: NIH Image to ImageJ: 25 years of image analysis. Nat Methods 2012, 9:671–675.

33. Miller JH: Experiments in molecular genetics. Cold Spring Harbor Laboratory; 1972.

34. Karimova G, Pidoux J, Ullmann A, Ladant D: A bacterial two-hybrid system based on a reconstituted signal transduction pathway. Proc Natl Acad Sci 1998, 95:5752–5756.

35. Lori C, Kaczmarczyk A, de Jong I, Jenal U: A Single-Domain Response Regulator Functions as an Integrating Hub To Coordinate General Stress Response and Development in Alphaproteobacteria. mBio 2018, 9.

36. Li S, Yu X, Beattie GA: Glycine Betaine Catabolism Contributes to Pseudomonas syringae Tolerance to Hyperosmotic Stress by Relieving Betaine-Mediated Suppression of Compatible Solute Synthesis. J Bacteriol 2013, 195:2415–2423.

## References

1. Martínez-García E, de Lorenzo V: Engineering multiple genomic deletions in Gram-negative bacteria: analysis of the multi-resistant antibiotic profile of Pseudomonas putida KT2440: Tools for editing Gram-negative genomes. Environ Microbiol 2011, 13:2702–2716.

2. Miller WG, Leveau JHJ, Lindow SE: Improved gfp and inaZ Broad-Host-Range Promoter-Probe Vectors. Mol Plant-Microbe Interactions® 2000, 13:1243–1250.

3. Bao Y, Lies DP, Fu H, Roberts GP: An improved Tn7-based system for the single-copy insertion of cloned genes into chromosomes of gram-negative bacteria. Gene 1991, 109:167–168.

4. Mossessova E, Lima CD: Ulp1-SUMO Crystal Structure and Genetic Analysis Reveal Conserved Interactions and a Regulatory Element Essential for Cell Growth in Yeast. Mol Cell 2000, 5:865– 876.

5. Sambrook J, Russell DW: Molecular cloning: a laboratory manual. Cold Spring Harbor Laboratory Press; 2001.

6. Stutz EW: Naturally Occurring Fluorescent Pseudomonads Involved in Suppression of Black Root Rot of Tobacco. Phytopathology 1986, 76:181.

7. Regenhardt D, Heuer H, Heim S, Fernandez DU, Strompl C, Moore ERB, Timmis KN: Pedigree and taxonomic credentials of Pseudomonas putida strain KT2440. Environ Microbiol 2002, 4:912–915.

